# The pursuit of motivational goals reduces pain through an opioidergic mechanism

**DOI:** 10.64898/2026.08.26.747254

**Authors:** Leonard Asan, Ole Goltermann, Lars Keuter, Jonathan Jeßberger, Christian Büchel

## Abstract

Pain promotes protective behavior but can interfere with other biologically important goals. Survival may require overcoming pain to obtain rewards, secure resources or escape danger, yet evidence for pain modulation by competing demands and endogenous modulatory systems during goal pursuit is lacking. We developed a paradigm in which participants chose whether to pursue monetary rewards despite painful heat stimulation during fMRI, under placebo or opioid receptor blockade with naloxone. Actively pursuing motivational goals during painful stimulation reduced perceived pain and increased fMRI signal in pain-modulatory cortical regions, including multiple subregions of the rostral anterior cingulate cortex (rACC) and dorsolateral prefrontal cortex, alongside enhanced rACC-periaqueductal gray coupling, consistent with recruitment of the descending pain modulatory system. Behavioral and neural effects were attenuated by naloxone, supporting a mediating role for endogenous opioids. These findings provide convergent evidence that active goal pursuit engages opioidergic pain modulatory mechanisms to reduce pain in humans.

## Introduction

Pain serves a core motivational function: it signals tissue damage and motivates the organism to engage in nocifensive responses to prevent or limit harm (*1*). This has obvious evolutionary advantages, as absence of nocifensive behaviors may worsen damage and impair healing. However, survival often requires behaviors which conflict with nocifensive responses, such as foraging despite injury during scarcity or escaping a predator after being harmed (*2*). In these situations, acute pain conflicts with the pursuit of important goals. One possibility to solve this conflict is a motivation-decision arbitration (*3*, *4*), which suggests that when a noxious stimulus occurs in the presence of a competing motivational goal, the agent makes an implicit decision about which of the behavioral responses is more important. If the behavioral response necessary to obtain the competing motivation-driven goal is deemed more important, pain is reduced. Previous observations of altered decision-making induced by the threat of pain (*5*), altered reward processing under pain (*6*), increased distraction-induced hypoalgesia for pain during rewarded vs. non-rewarded tasks (*7*) and hypoalgesic effects of reward presentation (*8–10*) have provided some indirect evidence for such a mechanism. However, these studies focused on changes of pain-related behaviors, rather than changes in pain itself, or on how reward outcome, rather than a motivation-driven goal such as reward pursuit, modulates pain. This is fundamentally distinct from the central prediction of a motivation-decision model, namely that pain reduction should arise directly from an organism’s decision to pursue a behaviorally relevant goal.

It was further suggested that the hypoalgesic effect associated with motivated decisions is mediated by a descending pain modulatory system (DPMS) (*3*, *4*). This network exerts top-down control from higher-order cortical structures, most importantly the rostral anterior cingulate cortex (rACC), via the midbrain periaqueductal gray (PAG) and rostral ventromedial medulla (RVM), to the dorsal horn of the spinal cord, employing endogenous opioids as an important neurotransmitter (*11–14*). This system is important for various psychological pain-modulatory effects such as placebo hypoalgesia (*15–17*), stress-induced hypoalgesia (*18*, *19*), distraction-related hypoalgesia (*20*, *21*), or conditioned analgesia (*22*, *23*). However, no study to date has demonstrated opioidergic involvement in descending pain control triggered by the decision to pursue a motivational goal.

The present work was designed to provide convergent evidence for the core behavioral and neurobiological assumptions of such a mechanism. In a randomized, placebo-controlled experiment (Fig. 1), we tested the central predictions of such a model. To this end, we developed a novel task (Fig. 2), which participants completed while undergoing blood-oxygen-level-dependent (BOLD) functional magnetic resonance imaging (fMRI) on two separate days: once under a saline (SAL) placebo control and once under the opioid receptor antagonist naloxone (NLX). During the task, participants chose whether to pursue a motivational goal (i.e., monetary reward), by accepting or declining to engage in a behavioral action (i.e., exerting physical effort via a grip force dynamometer) despite enduring heat pain. As outlined in the preregistration (for details see Methods), we formulated the following hypotheses: (1) pain ratings are reduced in accept relative to decline trials; (2) the extent of the pain modulation depends on the magnitude of the monetary offers; (3) rACC and PAG activation during pain is modulated in accept compared to decline trials; (4) rACC-PAG functional connectivity is increased in accept versus decline trials; and (5) all of these effects are blunted under the opioid antagonist naloxone.

**Fig. 1.**
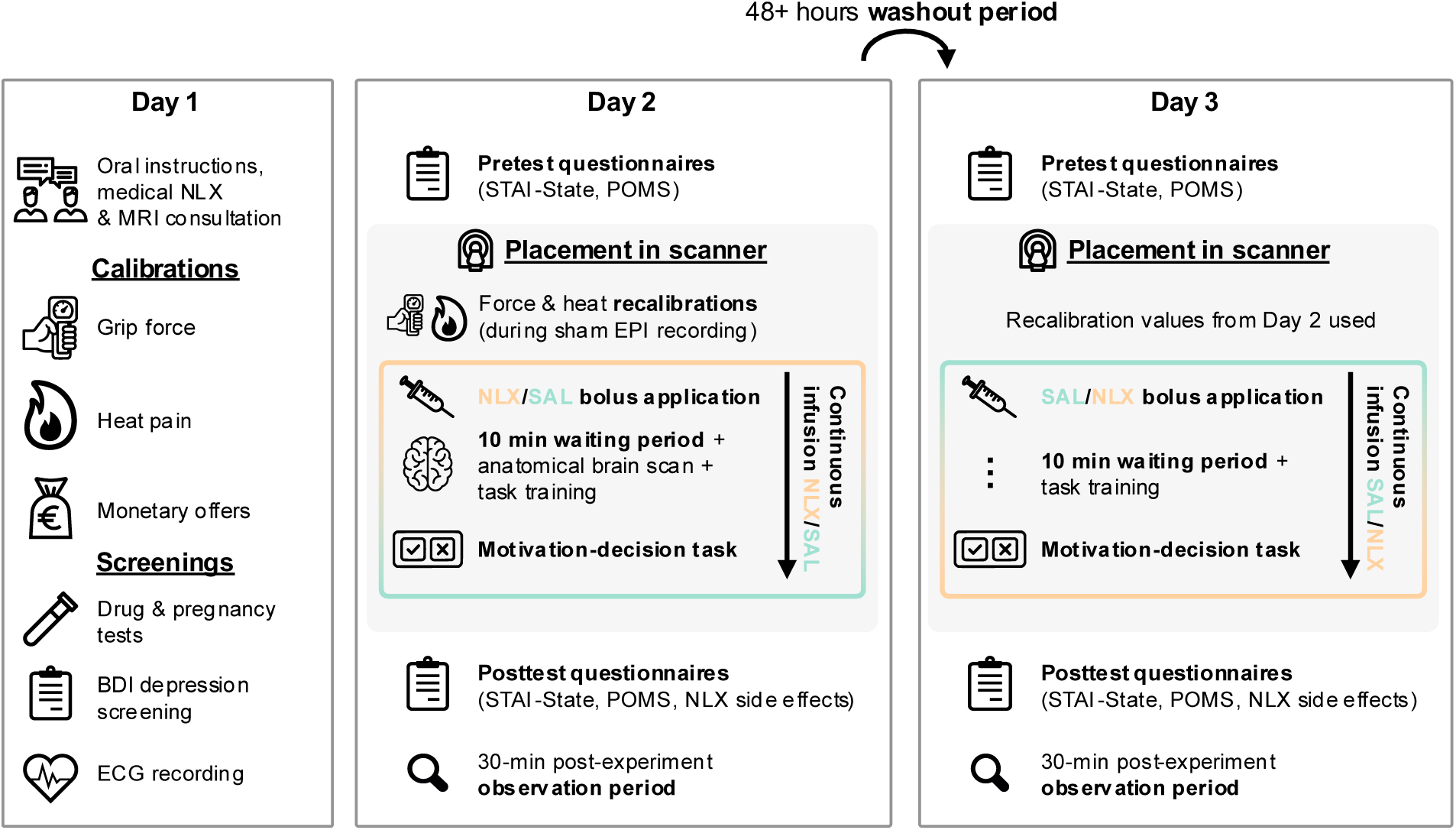
Experimental protocol. The study comprised three sessions. On day 1, participants received standardized instructions and a medical consultation regarding naloxone (NLX) administration and MRI safety. Individual calibrations were performed for maximal grip force, noxious heat stimulation, and monetary offers, followed by urine drug and pregnancy screening and an ECG recording. On days 2 and 3 (fMRI sessions), participants first completed questionnaires assessing state anxiety and mood. Inside the scanner, grip force and heat calibrations were repeated on day 2 only; calibration values from day 2 were used on day 3. Participants then received an intravenous bolus of NLX (0.15 mg/kg) or saline (SAL), followed by continuous infusion (0.2 mg/kg/h). During the 10-minute phase until a steady-state drug concentration was reached a T1-weighted scan was acquired and four practice trials of the motivation–decision task were performed (Fig. 2). Post-scan, questionnaires assessing state anxiety, mood, and NLX side effects were completed during a 30-min observation period. The third session, conducted after a washout period of at least 48 hours, replicated the fMRI session with the alternate drug condition (without recalibrations or T1 acquisition). Each fMRI session lasted approximately 2.5 hours in total, including approximately 1.5 hours in the scanner. POMS, Profile of Mood States; STAI, State-Trait-Anxiety Inventory.

**Fig. 2.**
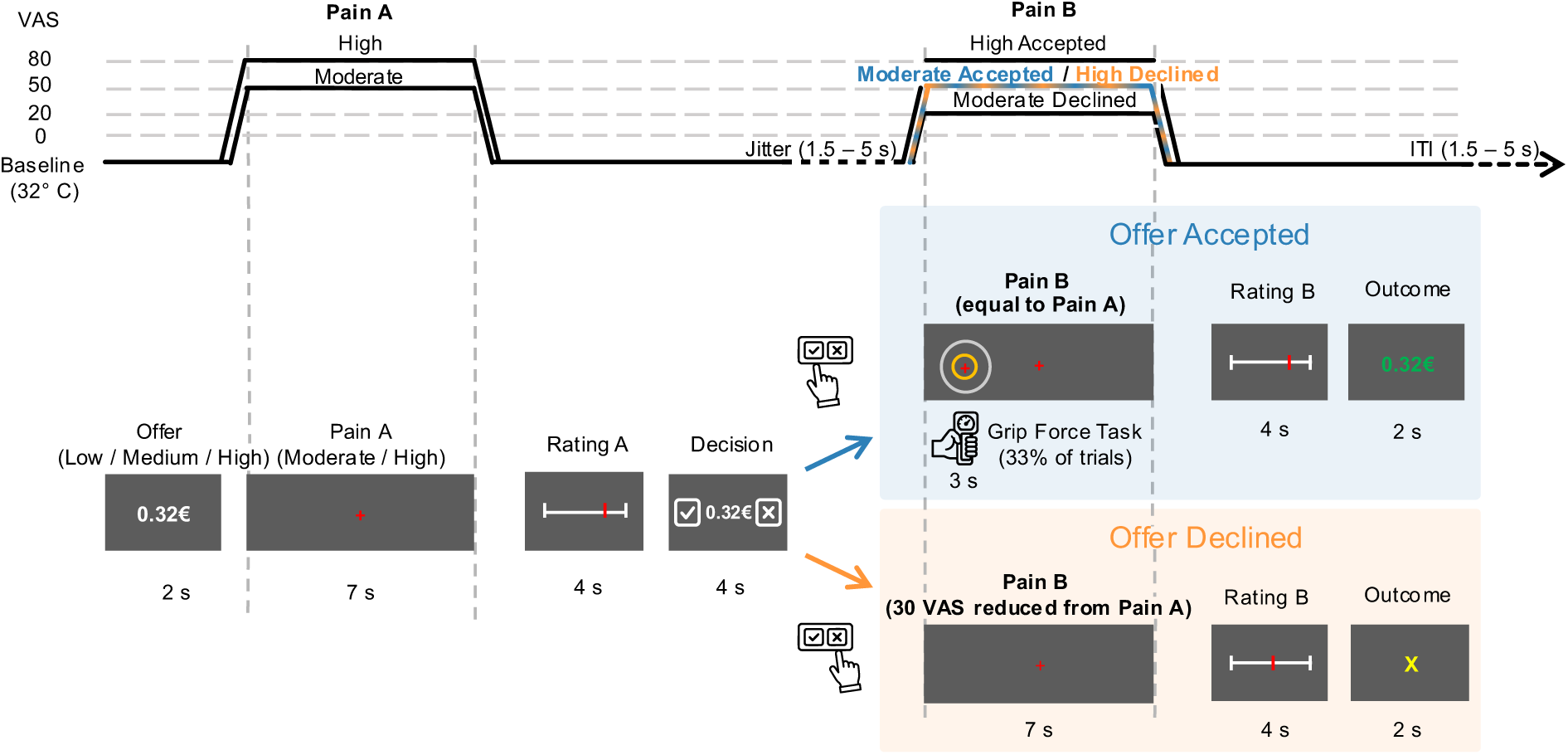
Schematic of a trial of the motivation-decision task. Each trial began with presentation of a monetary offer (low, medium, or high), followed by a 7-second painful stimulus (Pain A; calibrated to moderate or high intensity). Participants rated the pain intensity of the stimulus on a visual analog scale (VAS) before deciding whether to accept or decline the offer. Accepting resulted in a second painful stimulus (Pain B) delivered at the same intensity as Pain A, whereas declining reduced the intensity by 30 VAS units. This design enabled comparison of physically identical Pain B stimuli following accepted versus declined offers (blue and orange time courses). In one third of accepted trials (randomized), participants performed a grip-force task (grip force at 30% of individual maximum) during the first three seconds of Pain B. This task operationalized the physical response competing with the nocifensive response in the motivation-decision framework (see Methods). After Pain B, participants again rated pain, followed by trial outcome feedback. Each session comprised 72 trials.

## Results

### Effective pain induction

As an initial validation step, we confirmed successful induction of heat pain by examining both behavioral ratings and BOLD responses to Pain A (pre-decision). Participants rated high-intensity heat stimuli as significantly more painful than moderate-intensity stimuli (*t*(52.7) = 18.96, *p* = 3.36 × 10^−25^; Supplementary Fig. S1a), confirming successful pain induction. Blocking opioid receptors with NLX led to an overall increase in pain perception (main effect of drug: *F*(1, 6599.4) = 28.53, *p* = 9.56 × 10^−8^) and a significant drug-by-intensity interaction (*F*(1, 6634.1) = 11.61, *p* = 6.59 × 10^−4^). NLX enhanced pain perception more strongly at moderate intensity (mean difference = 3.68 VAS) than at high intensity (mean difference = 0.83 VAS). In the fMRI data, high (vs. moderate) pain elicited robust, whole-brain family-wise error (FWE) corrected activations across drug conditions in canonical pain-processing regions, including the anterior insula (xyz in mm: 36, 8, 8; *Z >* 8, *p* < .001), dorsal posterior insula (38, –15, 18; *Z* > 8, *p* < .001), mid-cingulate cortex (–9, 14, 36; *Z* > 8, *p* < .001), primary somatosensory cortex (26, −30, 62; *Z* = 7.99, *p* < .001), secondary somatosensory cortex/parietal operculum (50, −27, 27, *Z* > 8, p < .001), and thalamus (14, –14, 2; *Z* = 7.77, *p* < .001; Supplementary Fig. S1b). Increased activity during pain across intensities under NLX relative to SAL was observed in thalamus and cerebellum (Supplementary Table 1). No voxels survived whole-brain correction for the drug-by-intensity interaction contrasts. Together, the results confirm successful pain induction and activation of pain-sensitive brain areas.

### Decision-related effects on pain processing

#### Pain perception before the decision predicts subsequent decision-making

Across intensities, trials associated with lower Pain A ratings were more likely to be followed by acceptance of the offer (mean difference = 9.26 VAS, *SE* = 1.05, *t*(39.5) = 8.84, *p* = 6.68 × 10^−11^; Supplementary Fig. S2). There was no evidence for an interaction between decision and intensity (*p* = .214), drug condition and decision (*p* = .162), or a drug-by-decision-by-intensity interaction (*p* = .081). Results for Pain A confirm an association between pain intensity and subsequent decision during motivational conflict.

#### Opioidergic pain reduction following active accept decisions

After establishing an association between pain and decisions for Pain A, we next tested the central prediction of a motivation-decision mechanism (*3*), namely that pain is attenuated after deciding to pursue a motivation-driven goal, relative to pain for an identical stimulus after declining to engage. To this end, we analyzed ratings for pain after the decision (Pain B), focusing on trials in which participants declined an offer following high-intensity pain, and trials in which they accepted an offer following moderate-intensity pain. Critically, in both conditions, Pain B was of the same intensity (50 VAS). This allowed us to compare pain perception for physically identical stimuli. Furthermore, we included per-trial Pain A ratings (z-standardized within participant, session, and stimulus intensity), together with their interactions with decision and drug condition, as covariates in the linear mixed-effects model (see Methods for full model specification). This approach controlled for trial-by-trial variability in pain sensitivity, allowing us to isolate the effect of the decision on subsequent Pain B ratings beyond momentary pain fluctuations.

In accordance with our preregistered hypothesis, pain ratings were lower following accepted compared to declined offers (main effect of decision: *F*(1, 51.9) = 14.09, *p* = 4.41 × 10^−4^). Crucially, as hypothesized, blocking opioid receptors with NLX blunted this effect (decision-by-drug interaction: *F*(1, 3900.4) = 10.28, *p* = .001; Fig. 3), as the effect of decision was substantially larger in the SAL condition (mean difference declined vs. accepted = 5.66 VAS, *SE* = 1.29, *t*(66.4) = 4.38, *p* = 4.22 × 10^−5^) than under NLX (mean difference declined vs. accepted = 2.98 VAS, *SE* = 1.27, *t*(62.5) = 2.34, *p* = .023). Together, these findings provide evidence that in humans, the decision to pursue a motivational goal leads to pain reduction, and that this reduction in pain involves opioid receptor signaling.

**Fig. 3.**
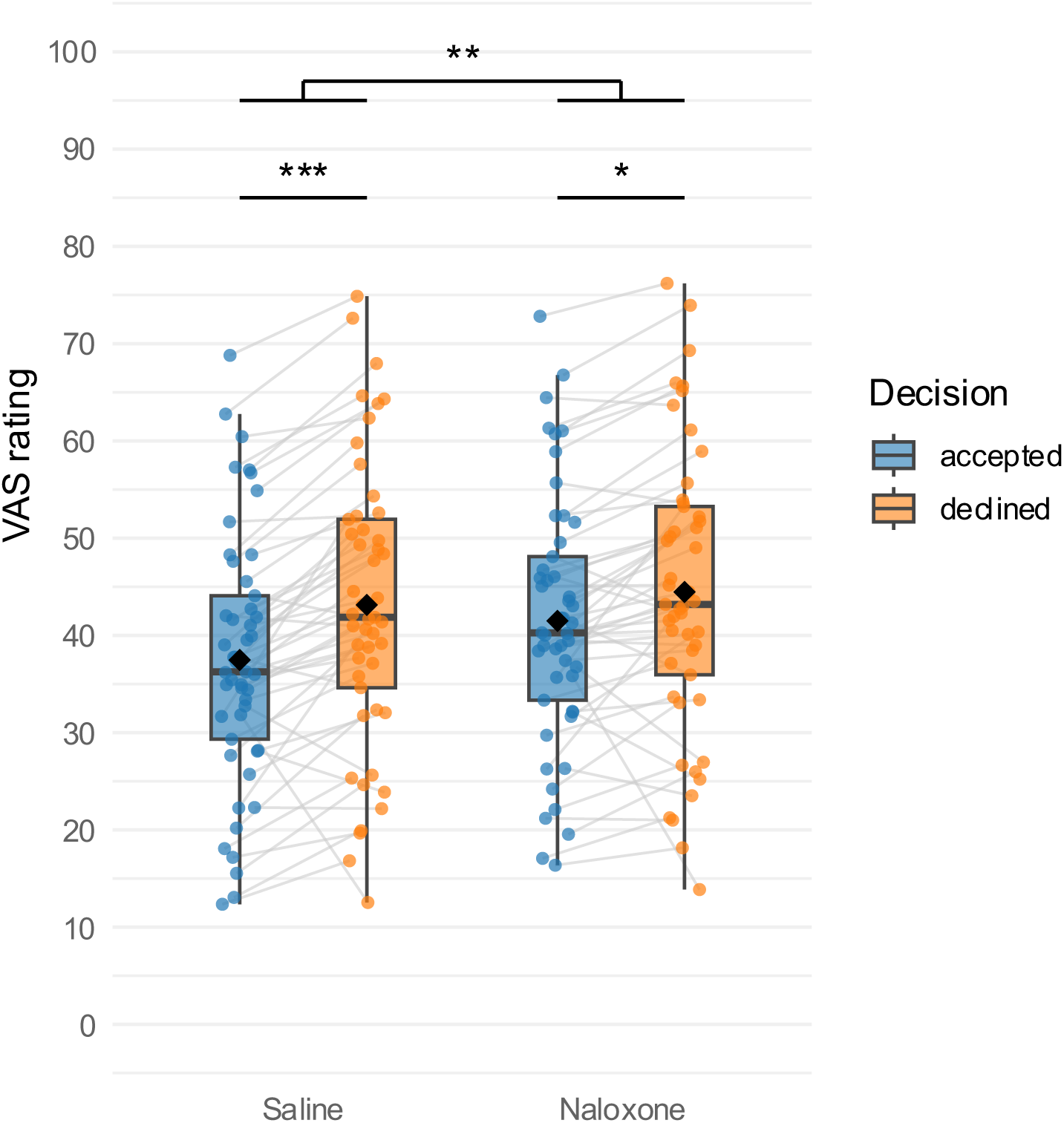
Decisions to pursue a motivational goal reduce perceived pain. Pain after accepting an offer was decreased relative to pain after declining an offer. The modulatory effect of decision on pain was reduced by NLX, as reflected in a significant decision-by-drug interaction across offer levels, indicating an opioidergic mechanism. Diamonds indicate estimated marginal means, lines denote medians, individual data points denote participant-level estimated marginal means (including random effects). Gray lines indicate within-subject differences between accept and decline conditions. *N* = 49. VAS, visual analog scale. \**p* < .05, \*\**p* < .01, \*\*\**p* < .001.

Next, to confirm that the observed difference between Pain B after accepted vs. declined offers was not driven by the intensity of the preceding Pain A stimulus (50 VAS for accepted offers, 80 VAS for declined offers), we performed a control analysis across trials (see Supplementary Text 1: Control Analysis I, for details). Here, we compared 50 VAS Pain A stimuli which were preceded by either an 80- or 50 VAS Pain B in the previous trial. Because on average inter-stimulus intervals between pain stimuli were deliberately kept identical within and across trials, this comparison matched the decision-related contrast in all physical properties. No significant difference was observed (mean difference = 0.47 VAS, *p* = .497). Bayesian model comparison further supported the absence of a carry-over effect (Bayes factor BF_01_ = 3.80), indicating that the data were approximately 3.8 times more likely under the model excluding a carry-over effect than under the model including it. This confirms that the decision-related difference in pain perception is not driven by carry-over effects linked to the physical properties of the preceding pain stimulus. In addition, we observed a significant effect of the effort task which had to be performed in one third of all accept decisions (*F*(1, 3894.8) = 18.44, *p* = 1.80 × 10^−5^). Pain reduction was greater in trials in which participants executed the grip force task compared with trials without effort (mean difference = 1.96 VAS). Because the effort task occurred only on “accept” trials, we assessed whether the decision-related effects could be attributed primarily to properties of the grip force task, such as distraction, attentional load, motor- or exercise-induced hypoalgesia, or conditioned pain modulation resulting from prolonged or aversive isometric contractions (*20*, *21*, *24–27*). We minimized the effects of these confounds by keeping the grip force task short (3 seconds) and low in intensity (30% of maximal grip force). Nevertheless, to assess a possible residual influence of task effects, we repeated the Pain B analysis restricted to trials without the effort task (see Supplementary Text 1: Control Analysis II). This analysis reproduced the original pattern of results (Supplementary Fig. S3), demonstrating that the decision- and drug-related modulation of pain cannot be explained by task demands alone. In one respect, we deviated from our preregistration, as we could not assess the influence of offer amount on decision-related pain modulation, because the high acceptance rates for some offers (Supplementary Fig. S4) precluded reliable estimation of decision-by-offer interactions in the behavioral or fMRI data. A third preregistered behavioral analysis examined whether drug or offer modulated within-trial changes in perceived pain, quantified as the difference between Pain A and Pain B on “accept” trials. Results showed larger pain reduction from Pain A to Pain B in the SAL condition relative to the NLX condition (main effect of drug: *F*(1, 5101.6) = 5.25, *p* = .030). Medium (mean difference = 1.34 VAS, *p* = .002) and high (mean difference = 1.34, *p* = .002), but not low offers (mean difference = 0.78, *p* = .201) were associated with significant reductions from Pain A to Pain B (Supplementary Fig. S5). These effects should be interpreted with caution, because the monetary offer preceded both pain phases in every trial, and thus a modulatory influence could act on Pain A as well as Pain B. As a result, the difference between the two ratings does not cleanly isolate changes specific to motivation in the way the main analysis of Pain B ratings does.

#### Reward anticipation in the absence of self-determined goal pursuit does not predict pain reduction

Motivation comprises multiple interacting components, including reward anticipation, value attribution, and decision-making (*28*, *29*). Having established that pain is reduced after accepting an offer, an important next question concerns which aspect of motivation drives this analgesic effect. One potential factor is reward anticipation. In our design, accepting an offer reflected a motivated state directed toward obtaining a goal and thus necessarily entailed the anticipation of a potential reward, whereas declining an offer eliminated both goal pursuit and reward anticipation. In order to dissociate reward anticipation from goal pursuit following an active decision, we conducted a follow-up behavioral experiment (see Supplementary Text 2 for details). Participants were presented with cues signaling rewards, losses, or neutral outcomes, followed by a 7-s painful stimulus and subsequent outcome presentation. Critically, to isolate reward anticipation effects from self-generated motivational choice, we removed the decisional component and the effort task and set the outcome probability to 100% to induce reward expectation on every trial. We expected that if reward anticipation alone contributes meaningfully to hypoalgesia, pain should be reduced in trials with a reward cue compared to neutral cue and potentially loss cue trials. However, pain ratings did not differ between reward and neutral trials (*p* = .338) or between reward and loss trials (*p* = .867; Supplementary Fig. S6). Bayes factor analysis provided strong evidence for the absence of a reward anticipation cue effect: a reduced model excluding the cue factor was approximately 38 times more likely to generate the observed data than a model including a reward anticipation effect (BF_01_ = 38.36). Moreover, sensitivity analyses indicated that the control experiment had sufficient power to detect effects substantially smaller than the decision-related hypoalgesic effect observed under SAL in the main experiment (Supplementary Text 2, Results). Specifically, the sample size of *n* = 18 provided 90% power (α = 0.05, two-tailed) to detect an effect size of *d* = 0.29, whereas the decision effect in the main experiment was *d* = 0.49. Together, these findings suggest that reward anticipation alone is unlikely to account for the pain modulation observed in the main experiment. Instead, the hypoalgesic effect appeared to be linked to participants’ motivational choice: When controlling for offer level, pain ratings were reduced after deciding to engage in an action in pursuit of the goal, consistent with a motivation-decision mechanism.

#### Increased opioid-dependent rostral anterior cingulate cortex engagement following accept decisions

We next examined the neural mechanisms underlying the observed behavioral effects. Specifically, we tested the hypothesis that the decision-related effects are mediated by activation of the DPMS, particularly the rACC and PAG, and that they are reflected in enhanced functional connectivity between these regions during pain following accepted versus declined offers. In line with the behavioral analyses, we focused on neural responses to Pain B (post-decision). Analyses were restricted to trials with identical pain intensity (VAS 50) which were preceded by either an accept or a decline decision (Fig. 4). We excluded trials in which participants executed the grip force task to ensure comparability of pain stimuli between accept and decline decisions and to avoid motion-related fMRI signal changes. All *p*-values reported in the following are FWE-corrected (i.e. corrected for multiple comparisons) for the relevant regions of interest (see Materials and Methods). In the SAL condition, accepting relative to declining an offer was associated with larger activation during pain in several subregions of the rostral anterior cingulate cortex (left pregenual ACC [pgACC]: –14, 42, 16; *Z* = 4.23, *p* = .007; ventral perigenual ACC [pACC]: left, –12, 45, –6; *Z* = 3.99, *p* = .018; right, 6, 45, –3; *Z* = 3.70, *p* = .049). We next examined whether this activation reflected endogenous opioid signaling by testing the effect under NLX. Strikingly, the decision-related rACC activity during pain observed under SAL was absent in the NLX condition; no significant differences in activation within the a priori defined PAG/rACC mask were observed (all *p* values > .22). This attenuation was confirmed by a significant decision-by-drug interaction in the rACC (left pgACC, –14, 42, 16; *Z* = 4.48, *p* = .003; right ventral pACC, 8, 45, –2; *Z* = 3.74, *p* = .043). Descriptively, at the peak coordinate in the left pgACC, 71.4% of participants showed greater activation during pain following accepted relative to declined offers under SAL (34.1% under NLX). Among participants showing positive accept-related activation, 90% exhibited larger effects under SAL than under NLX. A similar pattern was observed in left ventral pACC, where 69% of participants showed greater activation for accepted versus declined trials under SAL (46.3% under NLX), with 75.9% of these participants displaying reduced activation under NLX. In right ventral pACC, 73.8% showed greater activation under SAL (43.9% under NLX), and 80.6% of participants with positive accept-related effects exhibited NLX-induced reductions in this effect. BOLD responses in the PAG did not differ significantly between the accept and decline conditions under SAL or NLX (*p* values > .478). Notably, beyond the rACC, an exploratory whole-brain analysis revealed whole-brain FWE-corrected increased signal in the left dorsolateral PFC (dlPFC) (xyz in mm: −24, 44, 30; *Z* = 4.92; *p* = .038; Supplementary Fig. S7). 76.2% of participants showed greater responses during pain after accepting relative to declining in the left dlPFC region (43.9% under NLX). Among these subjects, 71.9% showed higher responses under SAL relative to NLX. This response attenuation under NLX was substantial but showed only a trend at the whole-brain level (decision-by-drug interaction: −24, 45, 30; *Z* = 4.82, *p* = .058).

**Fig. 4.**
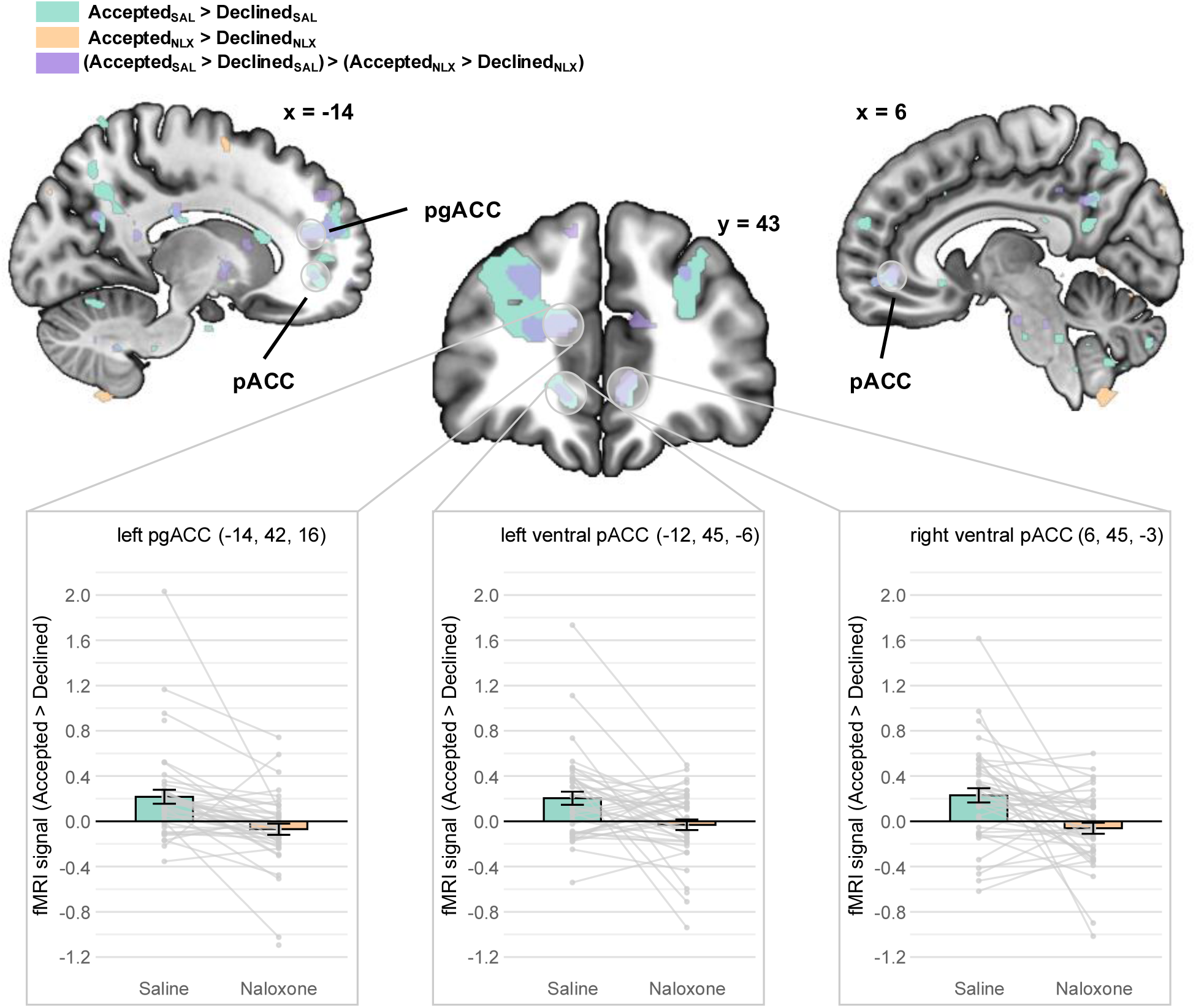
fMRI responses to Pain B (post-decision) in rACC. In the SAL condition, activation during accepted versus declined offers was increased in the left/ipsilateral pregenual ACC (pgACC) and bilateral ventral perigenual ACC (pACC). This activation was attenuated by NLX. The plotted parameter estimates were extracted from group-level peak voxels defined by the decision contrast under saline (Accepted_SAL_ > Declined_SAL_). Statistical maps are displayed at *p* < .001 uncorrected (for visualization only). Error bars represent standard errors of the mean. fMRI signal units are arbitrary. *N* = 45. ACC, anterior cingulate cortex; NLX, naloxone; pg, pregenual; SAL, saline; vp, ventral perigenual.

#### Increased opioidergic coupling between rACC and PAG after motivated decisions

After establishing involvement of the rACC in pain modulation after motivational conflict, we next examined whether activity in the rACC modulates the descending pain control pathway as a consequence of the motivated decision. Using participant-specific peak activation from the “Accepted > Declined” contrast in the SAL condition as a seed region, we tested functional connectivity of the rACC (Fig. 5). In line with our preregistered hypothesis, we observed stronger coupling between the pregenual ACC (pgACC) and the periaqueductal gray (PAG; xyz in mm: −3, −28, −9; *Z* = 3.58; *p* = .007) during accepted relative to declined trials, consistent with increased recruitment of the DPMS during goal pursuit. Again, we assessed whether this effect was mediated by endogenous opioids by repeating the analysis under NLX. As predicted, blocking opioid receptors with NLX abolished this coupling (all *p* values > .628), reflected in a significant decision-by-drug interaction within the same PAG cluster (−3, −30, −9; *Z* = 4.02; *p* = .001). At the reported PAG peak, 69% of participants showed increased functional coupling with the left pgACC under SAL during pain following accepted relative to declined offers (41.5% under NLX). Among participants showing positive coupling under SAL, 82.8% exhibited reduced connectivity under NLX. No significant changes in connectivity with the PAG were observed for ventral pACC bilaterally, suggesting a specific role of the pgACC in mediating a descending pain modulatory effect related to pursuing a motivational goal.

**Fig. 5.**
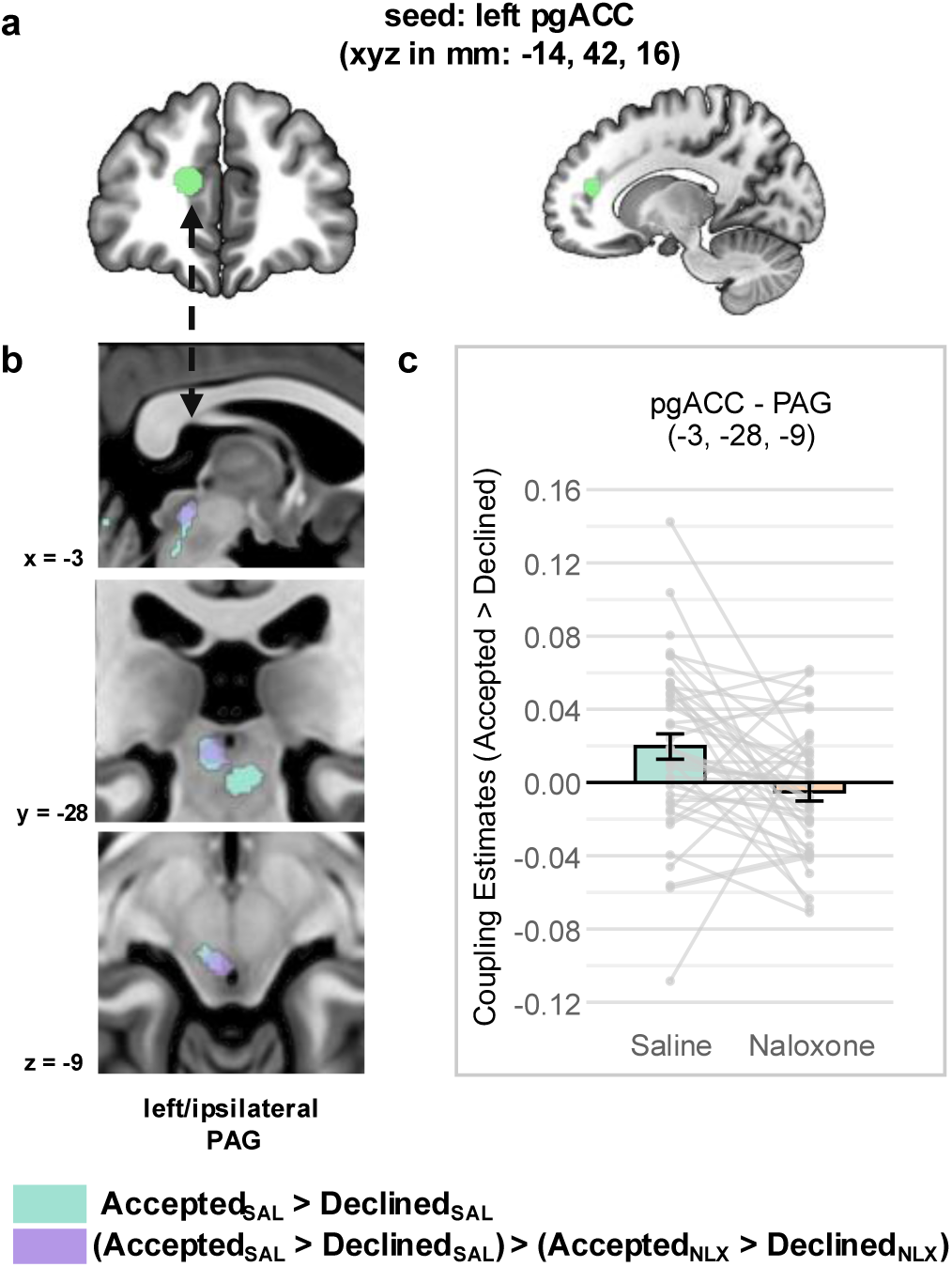
Functional connectivity results. **(a)** Seeds were defined based on the subject-specific peak activation within a 6-mm-radius sphere around the group-level peak activation in the left pgACC from the contrast Accepted > Declined under SAL. **(b)** The analysis revealed increased functional connectivity between the left pgACC and the left PAG during pain following accepted compared to declined offers under SAL. This effect was not observed under NLX, as indicated by a significant decision-by-drug interaction. Threshold for visual display was set at *p* < .001 uncorrected. **(c)** Parameter estimates visualizing differential connectivity strength for Accepted > Declined decisions in the SAL and NLX conditions. Points and lines represent individual subject values. Error bars represent standard errors of the mean. Coupling estimate units are arbitrary. *N* = 45. ACC, anterior cingulate cortex; NLX, naloxone; PAG, periaqueductal gray; pg, pregenual; SAL, saline.

## Discussion

Our data provide evidence for hypoalgesia during the pursuit of motivational goals in humans. The convergent behavioral, pharmacological and fMRI findings are consistent with the involvement of opioidergic descending pain-modulatory mechanisms proposed by a motivation-decision model of pain regulation (*3*). Specifically, we found increased activation during pain after accepting relative to declining a monetary offer in the rACC. The pregenual rACC additionally showed increased functional connectivity with the PAG, a region which has repeatedly been shown to play an important role in descending opioid-mediated analgesia (*11*, *30*, *31*). This anatomical specificity is key to interpreting the findings within the motivation-decision framework. The pgACC has consistently been implicated in representing the value of rewarding outcomes and in anticipatory, motivational action-outcome processing (*32*, *33*). The identified PAG cluster corresponds to a key node within the circuitry supporting descending, opioid-mediated analgesia. The finding of specific pgACC-PAG coupling may indicate that the valuation of the chosen goal, represented in the pgACC activity, may directly drive pain modulation. Critically, both the increased rACC activation and the increased connectivity between the rACC and the PAG were absent under NLX, providing evidence for the involvement of endogenous opioids in these mechanisms.

In this study, NLX significantly reduced decision-related differences not only in regions implicated in downstream descending modulation, but broadly across the rACC. While studies have repeatedly shown opioid dependence of descending pain pathways downstream of the rACC (*15*, *31*, *34*), opioids have also been proposed to alleviate the affective, aversive component of pain (*35–37*) and modulate hedonic processing of outcomes (*38*) through signaling within the rACC. Thus, additional supraspinal opioidergic mechanisms may have contributed to the motivational pain modulation we observed. Supporting this notion, we found involvement of a region in the dlPFC which showed a trend for attenuated responses under NLX, consistent with prior work implicating the dlPFC in pain modulation (*39*, *40*), including a previous PET study assessing opioidergic involvement (*41*). The observed bilateral ventral perigenual ACC responses, which did not show increased motivation-related coupling with the PAG, may also relate more strongly to supraspinal cognitive-affective pain regulatory processes than to DPMS recruitment (*42*). It is further worth noting that while our results from the NLX blockade provide evidence for opioid involvement, the observed behavioral modulation was not entirely blocked by NLX. This suggests the possibility of contributions from other neuromodulators, such as endocannabinoids, which also play an important role in pain regulation and interact with the DPMS (*42–44*).

While we observed significantly increased functional connectivity between the pgACC and PAG under SAL, we did not detect PAG BOLD signal differences for the corresponding decision contrast at our specified threshold. Given the functional heterogeneity of the PAG, which extends beyond hypoalgesia (*45–47*), PAG signal of comparable magnitude during the decline condition may reflect distinct defensive or aversive processes. Importantly, activation and connectivity analyses capture distinct aspects of neural function: whereas the former tests condition-dependent differences in BOLD responses averaged over the entire stimulus duration, the latter assesses condition-dependent covariation between pgACC and PAG timeseries, independent of their mean response levels. Thus, altered pgACC-PAG coupling can occur in the absence of mean differences in regional PAG activity.

Our sample generally showed a high willingness to accept offers. Although the individual monetary calibration (see Materials and Methods) mitigated this tendency to some extent, it resulted in comparatively low offer magnitudes (Supplementary Fig. S8) and precluded the possibility of examining parametric effects of reward amount due to the low number of decline observations for some offer levels (Supplementary Fig. S4). Thus, while the decision-related modulation was robust, the extent to which it scales with an offer remains an open question. Participants reported slightly stronger NLX-related side effects during the NLX session (*p* = .037) and identified the pharmacological condition above chance level (25.5% above chance level). However, neither state anxiety (*p* = .084) nor mood (*p* = .340) differed significantly between drug conditions in pre-post session assessments (see Materials and Methods). Although all participants received medical consultation about NLX, they were not informed about its specific role in pain modulation or its function within the context of this study. Moreover, the direction of the study hypotheses and the planned comparisons were not disclosed. Thus, while some participants may have been aware of the substance administered, such awareness is unlikely to have systematically biased the experimental outcomes.

Goal pursuit is inherently multifaceted. Committing to a goal following a decision entails multiple interrelated processes, including agency, goal commitment, action preparation and initiation (*48*, *49*). These constituent aspects may jointly shape the influence of goal pursuit on pain. Because pain modulation during goal pursuit had not previously been demonstrated in humans, the present study focused on the effects of the integrated state arising from a decision to pursue a motivational goal. Future studies which selectively manipulate its individual components may help dissociate their respective contributions to pain processing.

Our findings have important theoretical and practical implications. They show that pain modulation can arise from internally generated motivational states rather than external suggestions or stressors. Unlike other psychological hypoalgesic effects for which DPMS recruitment has been shown, like placebo hypoalgesia (*15–17*), stress-induced hypoalgesia (*18*, *19*), distraction-related hypoalgesia (*20*, *21*), or conditioned analgesia (*22*, *23*), the decision to accept an offer in our task was self-initiated and instrumental - it directly determined whether participants would pursue or avoid an alternative goal. The resulting hypoalgesia thus reflects volitional engagement of descending control rather than externally generated expectancy, cognitive manipulation or other external circumstances.

The results further align with assumptions of the integrative psychobiological pain model (*50*), which unifies the motivation-decision model of pain with expectation, controllability, and learning-based accounts of pain persistence. This framework proposes that an arbitration process continually weighs the motivational relevance of pain against alternative goals, dynamically shaped by reward, agency, and expectation. When motivationally salient alternatives dominate, the model predicts DPMS-mediated pain reduction. Conversely, increased or prolonged pain experience over time biases the system toward negative expectations, fear of action, and perceived helplessness, shifting attention further toward pain and away from alternative motivational goals. Our findings empirically ground the motivation-decision component of this integrative model. Moreover, the active, self-generated modulation observed here may represent an opportunity to break the outlined vicious cycle. In line with this interpretation, strengthening motivational engagement may not only reduce pain acutely but also counteract the motivational and affective imbalances that contribute to chronicity and pain persistence. Our results thus deliver neurobiological evidence to substantiate the merits of motivation-based cognitive pain management therapies, for example in rehabilitation after surgery, or when coping with chronic pain (*51*).

## Materials and Methods

### Preregistration

The study was preregistered at the WHO-accredited German Clinical Trials Register (DRKS). It is available at https://drks.de/search/en/trial/DRKS00033920. The comprehensive preregistration protocol can be downloaded from the ‘Study protocol and other study documents’ section of the linked registry entry.

### Ethics

The protocol conformed to the standards laid out by the World Medical Association in the Declaration of Helsinki and was approved by the local Ethics Committee (Ethikkommission der Aerztekammer Hamburg, vote 2021-10501-BO-ff).

### Participants

An a priori power analysis (α = .05, power = 0.90) for a within-subjects repeated measures ANOVA indicated that a sample size of *N* = 49 participants was required to detect a behavioral condition-by-drug interaction effect corresponding to *d* = 0.60, with the predicted effect size derived from the condition-by-drug interaction reported in a previous naloxone study (*52*). In total, 67 participants were recruited for the study. Of the 67 recruited participants, 14 did not complete all three experimental sessions (see Supplementary Table 2 for a list of dropout reasons). For the first three of the remaining participants, stimulation temperatures for moderate and high pain intensities differed across the two MRI sessions due to pain recalibrations before the second session. We retained the first of these three participants, as stimulus temperature differences between sessions were negligible (less than .22°C difference within each intensity); data from the other two were excluded due to substantial differences. Subsequently, temperatures were kept constant between sessions. Additionally, two participants were excluded for insufficient pain application (over one third of moderate-intensity stimuli rated as 0 on a 0-100 VAS scale on at least one day). This resulted in a final sample of *N* = 49 participants (24 female; mean age = 25.55 years, *SD* = 4.71) included in all behavioral analyses. For fMRI analyses, four participants were excluded due to excessive head motion (n = 3: >6 mm maximum scan-to-scan head motion within runs; n = 1: >0.7 mm scan-to-scan motion in more than 5% of volumes in every run of one session), based on exclusion criteria previously applied in studies from our laboratory using comparable MRI acquisition and preprocessing parameters (*24*, *53*). Hence, the resulting sample for fMRI analyses included *N* = 45 subjects (23 female, mean age = 25.73, *SD* = 4.72). In the behavioral sample and fMRI subsets, 26 and 25 participants received NLX in the first experimental session, respectively. All participants were screened for the preregistered inclusion criteria listed in Supplementary Table 3, gave written informed consent prior to participation and were compensated for their participation.

### Experimental Protocol

The experimental protocol comprised three sessions on three separate days (Fig. 1). On the first day (pre-experimental calibration session), participants received standardized oral instructions detailing all three sessions. A resident physician then provided medical consultation regarding drug administration and MRI safety. Written informed consent was obtained prior to the start of the procedures. Participants subsequently completed calibrations for individual maximal grip force, noxious heat stimulation, and monetary offers to be presented during the experimental sessions. At the end of the calibration session, electrocardiogram (ECG) and blood pressure readings were recorded and reviewed by the physician, and a urine sample was collected for drug and pregnancy screening. Total session duration was approximately 1.5 hours. On the second day, participants completed pre-test questionnaires assessing state anxiety and mood. Afterwards, blood pressure was measured, and an attending physician or trained medical assistant inserted an intravenous line into the right antecubital fossa. Participants were then guided into the MRI scanner, where grip force and heat calibrations were repeated while EPI recordings were performed to mimic experimental conditions. Following recalibration, a bolus dose of 0.15mg/kg of either NLX or SAL was administered, followed by connection to a continuous infusion of 0.2mg/kg/h. A subsequent 10-minute waiting period ensured that a steady-state drug concentration was reached prior to task onset (*54*). During this period, a structural T1-weighted scan was acquired while participants completed four practice trials of the experimental task. Participants then performed the motivation-decision-task inside the scanner. Upon completion, feedback about the amount of money gained during the session was provided. Outside the scanner, participants were disconnected from the infusion, the IV line was removed, and blood pressure was measured. Post-experiment questionnaires assessing state anxiety, mood, and NLX-related side effects were completed during a 30-minute observation period, ensuring participant safety prior to discharge. The third session was completed after a washout period of at least 48h and was identical to the second, except that no recalibration of grip force or heat stimuli was conducted; parameters from the first fMRI session were reused. After completing the last session, participants guessed which session involved NLX administration, were debriefed about the experimental manipulation, and received monetary compensation. Total durations of session days two and three were approximately 2.5 hours each.

### Noxious heat stimulation

Thermal stimuli were delivered using a TSA-2 thermode (Medoc, Ramat Yishai, Israel) with a contact heat-evoked potential stimulator (CHEPS) applied to the volar surface of the left forearm. The baseline temperature was set to 32.0°C. For every noxious stimulus, temperature increased and decreased at a rate of 13°C/s, with a 7-second plateau maintained at target temperature. To prevent skin sensitization, the thermode position was changed after calibration and between runs. To this end, the volar surface of the left forearm was divided into a fixed calibration area and four sections corresponding to the four task blocks. Section order was randomized and counterbalanced across subjects and kept constant within participants for the two sessions. Following each repositioning, three low-intensity (10 VAS) preexposure stimuli were administered.

### Behavioral data acquisition

Computerized tasks for the motivation-decision paradigm, as well as calibrations of grip force, heat pain, and monetary offers, were implemented using Psychophysics Toolbox (*55*) version 3.0.18 for MATLAB (R2022a). Heart rate and respiration were recorded at 1,000 Hz using the Expression System (In Vivo, Gainesville, FL) for subsequent physiological noise modeling. Grip force was measured with an MRI-compatible hand clench dynamometer (BIOPAC Systems Inc., Goleta, CA) and acquired via AcqKnowledge (version 5.0, BIOPAC Systems Inc.). Real-time feedback of grip force was presented through data transfer from AcqKnowledge to MATLAB using BIOPAC’s Network Data Transfer (NDT) module. Pain ratings were provided using a visual analog scale (VAS; 0 = no pain, 100 = maximal endurable pain within the experimental context). Pain ratings and decisions were collected via a right-hand button box (HHSC-1×4-D, Current Designs Inc., Philadelphia, PA).

### MRI Data acquisition

Functional magnetic resonance imaging (fMRI) data were acquired using a 3 Tesla Siemens Magnetom PRISMA system (Siemens Healthcare, Erlangen, Germany) equipped with a 64-channel head coil. Functional images were obtained using a T2*-weighted echo-planar imaging (EPI) sequence with a multiband acceleration factor of 2 and additional in-plane acceleration (GRAPPA; factor 2). The sequence included 60 axial slices (2 mm slice thickness), a repetition time (TR) of 1.8 seconds, an echo time (TE) of 26 ms, and a flip angle of 70°. The field of view (FOV) was 224 by 224 mm and was positioned to include the upper medulla and brainstem. Voxel size was 2 mm isotropic. Structural images were acquired on the first experimental day (Day 2) using a T1-weighted MPRAGE sequence (1mm isotropic voxels, 240 slices). Shimming and auto-alignment took place at the beginning of each session before acquiring EPI images.

### Calibrations

#### Grip Force

To determine individual maximum grip strength, participants completed a standardized calibration procedure. A vertical on-screen scale provided real-time visual feedback. Participants were instructed to squeeze a force dynamometer with maximal effort and to maintain peak force as steadily as possible. A trial ended successfully once participants held their force steadily within a 9.8N (equivalent to 1 kg) window for one second. Each participant completed five successful trials. To reduce the influence of outliers, the highest and lowest values were discarded, and the mean of the remaining three trials was used as the participant’s calibrated maximum force.

#### Thermal Pain

The thermal calibration protocol was adapted from Horing et al. (*56*). To allow for skin initialization, participants were first exposed to three pre-exposure stimuli (42.0°C, 42.5°C, and 43.0°C). Pain thresholds were then estimated using a probabilistic tracking procedure (*57*): six thermal stimuli of varying intensities were applied sequentially, and participants indicated whether each was “painful” or “not painful.” The intensity of each subsequent stimulus was adapted based on the previous binary response. The final stimulus in the sequence was defined as the participant’s individual pain threshold. Following threshold determination, four stimuli were presented at +0.5°C, +1.0°C, +2.0°C, and +1.0°C relative to the threshold. Participants rated each stimulus using a 0 (not painful) –100 (maximal tolerable pain) VAS for subjective pain intensity. A linear regression model was fit to these four ratings to estimate the mapping between stimulus temperature and perceived pain. To ensure reliable sampling across the full VAS range, the scale was divided into five bins (0–20, 20–40, 40–60, 60–80, 80–100). If no ratings were yet recorded in a given bin, the model-predicted temperature corresponding to the bin’s midpoint (e.g., 30 VAS for the 20-40 bin) was applied, and the resulting rating was used to update the regression. This iterative process continued until each bin contained at least one data point. From the final model, we derived individualized stimulus temperatures corresponding to 10 (pre-exposure), 20 (low), 50 (moderate), and 80 (high) VAS. Temperatures were manually adjusted if the 80 VAS calibration temperature exceeded a safety limit of 49°C; in such cases, it was capped at this boundary. To ensure perceptual discriminability, a minimum spacing of 1.0 °C was required between the low-, moderate-, and high-intensity stimuli. If this criterion was not met, the 80-VAS temperature was anchored and the 50 and 20 VAS temperatures were adjusted downward to preserve this spacing. If the 10 VAS pre-exposure stimulus was below 42 °C, it was increased to 42°C. In cases where calibration produced implausibly steep stimulus-rating slopes, temperatures were manually adjusted, ensuring equal (minimum 1°C) spacing between the 20-, 50-, and 80-VAS stimuli while keeping the calibrated 80-VAS temperature fixed. While the aim of these adjustments was to place stimuli within low-, moderate-, and high-intensity temperature ranges, the key factor was that stimulus temperatures were held constant within each participant across sessions.

#### Monetary offers

To determine individually calibrated monetary offer values corresponding to low, medium, and high subjective value relative to moderate and high experimental pain, participants completed a behavioral calibration task during the pre-experimental session. This task was presented as the first part of the experiment instead of a calibration session to minimize transparency of the procedure. In each trial, participants viewed a visual bar indicating pain intensity (50% or 80% filled, corresponding to moderate or high pain, respectively) alongside a monetary offer. The offer was randomly drawn from 20 logarithmically spaced values ranging from €0.01 to €5. This logarithmic scaling was selected to provide finer resolution in the lower value range (*52*). Participants chose to either accept or decline the offer. Accepting triggered a 7-second painful stimulus at the indicated intensity, during which participants were required to perform the grip force task (detailed below). The task was included to ensure that the physical effort which would be required in the experiment was incorporated into the subjective valuation process. Following the stimulus, participants rated the perceived pain intensity. Successful completion of the grip force task resulted in reward feedback corresponding to the offer presented in each trial. Declining the offer led to neither pain nor reward. The post-decision interval before the next trial was matched across accept and decline conditions to eliminate strategic time advantages for participants when declining offers. Each of the 20 monetary values was presented once per pain level, yielding a total of 40 randomized trials (duration approximately 20 minutes). At the end of the experiment, 10 of the 40 trials were randomly selected and paid out.

Individual decisions were then fitted using logistic regression to derive offer values corresponding to acceptance rates of 10–20% (low), 45–55% (medium), and 80–90% (high). In the main experiment, values per trial were randomly sampled from a uniform distribution of the participant-specific calibrated ranges within each level. Participants who did not accept any offers during the monetary calibration would have been excluded after the calibration session; this situation did not occur in the present study. For cases where the calibration did not yield meaningful estimates, preregistered default value ranges were applied: €0.01–0.20 (low), €0.40– 0.60 (medium), and €0.80–1.00 (high). Average monetary amounts across all participants included in the final sample were as follows: €0.05-0.16 (low), €0.33-0.44 (medium), €0.66-0.85 (high). Average and individual calibrated offers are displayed in Supplementary Fig. S8.

### Drug administration

Drug administration followed protocols established in previous studies conducted at the University Medical Center Hamburg-Eppendorf (*15*, *24*). A licensed physician or trained medical assistant administered either naloxone (NLX) or saline (SAL) intravenously via a line inserted into the participant’s right antecubital fossa. Participants received a bolus of 0.15 mg/kg body weight of NLX (Naloxon-ratiopharm® 0.4 mg/ml, Ratiopharm, Ulm, Germany) or SAL (Isotone Kochsalz-Lösung 0.9%, Braun) while in the scanner, at least 10 minutes prior to the start of the main experimental paradigm to ensure steady-state drug concentrations of NLX (*54*). Immediately following the bolus, a continuous intravenous infusion was initiated at a rate of 0.2 mg/kg/h and maintained throughout the scanning session. The infusion was terminated upon completion of the paradigm. Participants and all experimenters including the administering physician and research staff remained blind to treatment condition for the entire duration of data collection. Unblinding of the participants occurred only after the experiment, conducted by a research assistant who had no other involvement in the experimental sessions. Before being debriefed, participants were asked to indicate, in a forced-choice format, which treatment they believed they had received on each session day. Participants identified the correct treatment in 75.5% of cases (correct: *n* = 37, incorrect: *n* = 12; χ²(1, *N* = 49) = 12.76, *p* = 3.55 × 10^−4^). To compare self-reported side effects following NLX versus SAL administration, we used a non-parametric Wilcoxon signed-rank test. Participants reported significantly more side effects following NLX (*M* = 6.12, *SD* = 5.67) compared to SAL (*M* = 4.56, *SD* = 4.88; *V* = 673.5, *p* = .037), with a small effect size (matched-pairs rank biserial correlation *r* = .36). To assess influence of drug administration on state anxiety and mood, we fitted separate linear mixed-effects models for each outcome variable. Models included fixed effects for time (pre- vs. post experimental session), treatment (NLX, SAL), and their interaction, as well as a random intercept for participant. We found no significant interactions between treatment and time for state anxiety (*F*(1, 132) = 3.04, *p* = .084) or mood (*F*(1, 138) = 0.92, *p* = .340), suggesting that NLX did not affect participants’ state anxiety or mood differently from SAL in a pre-post comparison.

### Tasks

#### Motivation-decision paradigm

Participants completed the experimental task (Fig. 2) on two separate days, each under a different pharmacological condition (NLX or SAL). Each session comprised 72 trials divided into four blocks of 18.

Each trial began with a 2-second presentation of a monetary offer (low, medium, or high, individually calibrated). One second thereafter, participants received a thermal pain stimulus (Pain A) to the forearm, administered at one of two individually calibrated intensities corresponding to 80 (high) or 50 (moderate) on a 0–100 VAS. Following Pain A, participants rated perceived pain intensity using a unidimensional VAS (0 = no pain, 100 = maximal tolerable pain) within a 4-second window. Participants were then presented with the initial monetary offer again and were given 4 seconds to decide whether to accept or reject it. After a variable jitter (1.5–5 seconds, randomized), a second thermal stimulus (Pain B) was administered. The intensity of Pain B was contingent on the participant’s decision: accepting the offer resulted in the same intensity as Pain A; declining it reduced the stimulus intensity (80 → 50 VAS or 50 → 20 VAS). Rather than explicitly explaining these precise contingencies linking decision outcomes to stimulus intensities, participants were instructed that accepting would result in a second pain stimulus “comparable in intensity” to the first, whereas declining would lead to a “noticeable reduction” in the subsequent intensity, and that failure to decide in time would result in a repetition of the initial stimulus and no reward. In one third of the accepted trials (randomly determined), Pain B was accompanied by a grip force task (detailed below). These trials were discarded from fMRI analyses. After cessation of Pain B, participants provided a second pain rating using the same VAS scale. A 2-second outcome screen ended each trial. Trials were separated by a randomized inter-trial interval (ITI) of 1.5-5 seconds. To reduce participants’ potential assumptions of spurious contingencies between offer value and pain intensity, the first trial of each block was designed to be incongruent (i.e., low offer with high Pain A or high offer with moderate Pain A). The pairing of offers and pain intensities was otherwise randomized and counterbalanced within each block.

#### Grip force task

In one-third of accepted trials within the motivation-decision paradigm, a grip force task was implemented at the onset of Pain B. During this task, participants used a hand dynamometer to manipulate a red circle on screen (*58*), aiming to expand it beyond the boundary of a larger gray circle (set to 30% of the participant’s maximum grip force, calibrated prior to the task).

Participants were instructed to maintain force above this threshold for at least 1.5 seconds (half of the 3-second task duration) without applying excessive force. While not explicitly stated, the upper boundary was set at 80% of maximum force. They were further told that a trial’s success would depend on how steadily the force was maintained above threshold, with the system tracking their force fluctuations and implicitly applying either a lenient, or, in rare trials, a very strict success criterion. In reality, success was determined solely by maintaining force above threshold and below the upper boundary, and successful trials were rewarded probabilistically at a 90% reinforcement rate.

Constructing the grip force task in this manner served several key experimental purposes. First, the task operationalized the competing behavioral response postulated by the motivation-decision theory (*3*). We reasoned that this step was necessary, as the theory suggests that the necessity for a goal-directed, active physical response engages descending pain modulatory systems by competing with the natural nocifensive response to pain. Second, by limiting exertion at a modest intensity (30%) and implementing a purported steadiness success criterion, we effectively minimized excessive scanner-related motion artifacts while maintaining face validity from the participant’s perspective. Third, the low physical and cognitive demands as well as the short duration of the task compared to the pain stimulus intended to reduce potential confounds from distraction-induced, task-related hypoalgesia (*7*, *20*) or conditioned pain modulation stemming from task-related fatigue or discomfort in the arm and hand (*26*, *27*, *59*). Fourth, by employing reward in a probabilistic manner, we dissociated pain modulation from guaranteed reward, limiting potential reward-related hypoalgesia effects shown previously (*8*, *9*). These design choices were intended to ensure that the observed reductions in pain would reflect preparatory mechanisms for goal-directed action, driven by the decision to pursue a motivational goal.

### Experimental Design

The experimental design incorporated five within-subject factors: offer level (low, medium, high), pain intensity (moderate [50 VAS], high [80 VAS]), decision (accept, decline), task type (effort, no effort), and drug (SAL, NLX). Factors were fully crossed with two exceptions: first, because the effort task was only asked to be performed in one third of accepted trials, factor task type was nested within “accept” trials. Second, as the decision factor was determined by participant behavior, the number of accept and decline trials varied across individuals. This variability was accommodated in all statistical models using linear mixed-effects approaches that included subject-level random effects and, where feasible, by-condition random slopes. Per participant, the order of the drug condition (SAL, NLX) was assigned to the session days according to a randomized, counterbalanced, and automatically generated blinding list. Participants and all experimenters who interacted with participants were blind to the drug condition.

### Questionnaires

During the initial calibration session (day 1), participants filled out the Beck Depression Inventory-II (BDI-II) (*60*). To assess potential drug-related changes in affective state, participants also completed the state scale of the STAI (STAI-S) (*61*) and a short form of the Profile of Mood States (POMS-SF) (*62*) immediately before and after both experimental sessions (days 2 and 3). Following each session, participants filled out a side effect questionnaire assessing seven common adverse symptoms associated with NLX administration (lethargy, dry mouth, dry skin, blurred vision, dizziness, headache, nausea) on a 5-point Likert-type scale ranging from 0 (“not at all”) to 4 (“very strong”).

### Statistical Analyses

#### Behavioral Analyses

Behavioral data were analyzed using R (version 4.2.1, R Core Team) within the RStudio environment (version 2024.12.1). Preprocessing of behavioral data included the following steps: (i) mean-centering of all continuous predictor variables added in the models (i.e., trial numbers within runs and sessions), (ii) exclusion of pain ratings of 0 and 100 (to avoid floor and ceiling effects), and (iii) exclusion of extreme rating outliers defined as over or under three times the interquartile range within participant, pain intensity, and session. Pain ratings from the first (Pain A) and second (Pain B) thermal stimuli were modeled as outcome variables in separate linear mixed-effects models (LMMs), which were analyzed using the lme4 package in R (*63*). Main effects and interactions were tested using Type III ANOVAs with Satterthwaite’s approximation for denominator degrees of freedom, as implemented in the lmerTest package (*64*). Post hoc comparisons of estimated marginal means were corrected for multiple testing using Tukey’s method. Results were considered statistically significant at a threshold of α = .05.

#### Behavioral Model Specification

In both Pain A and Pain B models, trial number within run, trial number within the session, and session day were included as covariates of no interest to account for potential habituation or sensitization effects after switching thermode position, across the experimental session, and across session days, respectively. We further added offer amount (low, medium, high) as a control covariate. For the Pain A model, fixed effects of interest included pain intensity (moderate vs. high), decision (accepted vs. declined), and drug condition (SAL vs. NLX), along with all combinations of interactions between these factors. The Pain B model was restricted to a subset of trials in which participants either accepted the offer following a moderate Pain A stimulus or declined the offer following a high Pain A stimulus. This selection enabled a direct comparison of identical-intensity (moderate) Pain B stimuli under different motivational contexts (after accepting vs. declining an offer). Consequently, pain intensity was not included as a factor in the Pain B model. Instead, to account for potentially confounding state-dependent fluctuations in pain perception, Pain A rating (z-scored within participant, intensity, and session) was included as a covariate, along with its interactions with decision and drug condition. Lastly, trial type (effort trial vs. no effort trial) was included to control for the effect of the effort task on pain perception.

Random-effects structures for both models were specified following the principle of including the maximal structure justified by the design (*65*). We thus considered participant-level random intercepts and random slopes for all fixed effects, except for session day and drug condition. These two factors were not modeled as random slopes because each participant contributed only one drug condition per session, with all other experimental manipulations nested within these sessions. As a result, drug condition and session were perfectly confounded at the within-subject level, precluding the estimation of independent variance components within participants for these factors.

To determine the most appropriate random-effects structure for our linear mixed-effects models fulfilling these criteria, we employed an automated approach using the buildmer package in R (*66*) with the argument [direction = “order”]. This procedure incrementally builds up the random-effects structure from a user-defined space of random intercepts and slopes, prioritizing model complexity without producing singular fits or convergence issues.

The final LMM specifications were as follows:

Pain A:

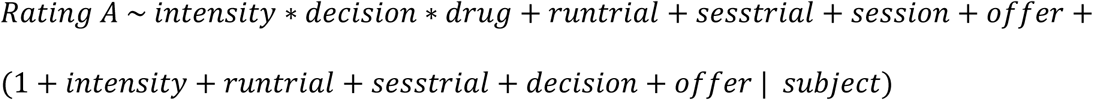

Pain B:

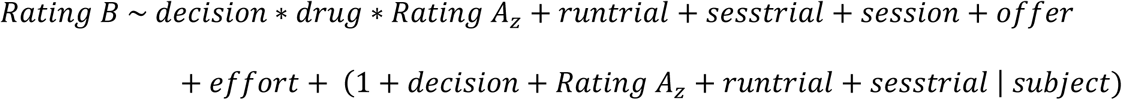

Where runtrial is the trial number within a run, sesstrial is the trial number within a session, and Rating A_z_ is the rating for Pain A within each trial, z-scored within participant, intensity, and session, offer is the offer amount (low, medium, or high; categorical) and effort is task type (categorical: effort trial vs. no effort trial).

#### Analysis of reductions in pain between Pain A and Pain B

To assess whether Pain B ratings following acceptance were reduced relative to Pain A, we fit an additional linear mixed-effects model using the difference between Pain A and Pain B ratings as the outcome, restricting the analysis to trials in which participants accepted the offer and thus received the same intensity during Pain A and Pain B. Contrary to our preregistered plan, we included both moderate and high intensity trials, as the final dataset contained sufficient numbers of “accept” decisions across all offer and intensity conditions to permit a reliable assessment of offer-related effects for this contrast. Consistent with the primary and control analyses of Pain A and Pain B, we specified the maximally justified random-effects structure for the within-subject design using the buildmer package (*66*).

The resulting model specification was as follows:

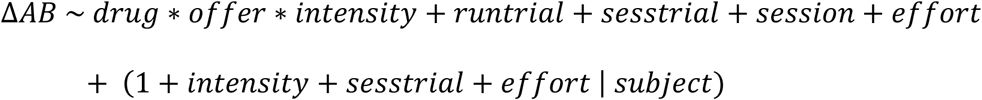

where ΔAB is defined as the difference between Pain A and Pain B ratings (Pain A - Pain B) in each “accept” trial. These effects should be interpreted with caution, because the monetary offer (i.e. the experimental manipulation) preceded both pain phases on every trial. Modulatory influence would therefore affect Pain A as well as Pain B, meaning that the difference between the two ratings does not cleanly isolate motivation-specific changes, unlike the main analyses.

#### MRI analyses

All MRI data were analyzed using SPM12 (Wellcome Trust Centre for Neuroimaging, London, UK) and MATLAB R2021b (The MathWorks Inc.). To ensure reproducibility and data accessibility, data were processed in BIDS format (*67*) using a standardized SPM12-based pipeline (available at [https://github.com/ChristianBuechel/spm_bids_pipeline.git]), which was used in previously published work (*24*, *53*). Figures displaying MRI data were generated using MRIcroGL (*68*). For visualization, all statistical maps were overlaid on the standard MNI152 template (mni152) included with MRIcroGL. All activation maps are shown at *p* < .001 uncorrected for display purposes, independent of the thresholds used for statistical inference.

#### Preprocessing of MRI data

Functional images from both experimental sessions were preprocessed jointly. Images first underwent slice timing correction and a first rigid-body realignment (six degrees of freedom). A non-linear coregistration step as implemented in the CAT12 toolbox (*69*) was then performed, aligning the mean EPI image from the first realignment step to the individual T1 image.

Coregistration with the T1 image was performed using the mean EPI generated during the first realignment step. The T1 image was normalized to MNI space (MNI152NLin2009cAsym) using DARTEL. Transformation fields were computed by combining the deformation field from the non-linear coregistration with the DARTEL flow fields, which were used to warp the individual EPI images into T1 and template space. A first-level analysis brain mask was created using gray and white matter tissue segments from the T1 image, warped into EPI space and slightly expanded (smoothed using a 3 mm FWHM Gaussian kernel). Finally, realignment was repeated, restricted to an individually generated brain mask to minimize the influence of non-brain related changes such as eye movements.

#### First-level GLMs

All single-subject general linear model (GLM) estimations were performed in native space on unsmoothed images. Per subject, the two sessions (SAL and NLX) were modeled in a joint GLM comprising a total of 8 runs. Constants for each run and each session were included in the models as covariates. All task regressors were convolved with a canonical hemodynamic response function (HRF). The primary regressors of interest modeled the two pain phases (Pain A and Pain B) as 7-second boxcars time-locked to stimulus plateau onset. Each pain phase was divided into separate regressors by the pain intensities existing for this phase (50 and 80 VAS for Pain A, 20, 50, and 80 VAS for Pain B), and further by decision outcome (accept vs. decline). For Pain B, regressors for pain during accepted trials were additionally split by whether an effort task was performed. In sum, 10 pain regressors were modeled per pharmacological condition (20 total).

Additional task regressors included 2-second boxcars modeling the offer and outcome phases, and a variable-duration boxcar for the decision phase (decision time + 1 second). Key presses were modeled as delta functions in a single regressor. A final task regressor modeled painful stimuli which were removed (i.e., those which were rated as 0 or 100, or which happened during trials in which the participant failed to make a decision). Nuisance regressors included 24 motion-related regressors (6 realignment parameters plus their squares and derivatives) (*70*) and 18 RETROICOR-based physiological noise regressors (*71*, *72*) generated using the physIO module of the TAPAS toolbox (*73*, *74*). To further account for physiological confounds, we included principal component regressors capturing signal from white matter (*n* = 6), cerebrospinal fluid (*n* = 6), and from a region around the posterior tip of the lateral ventricle (*n* = 6) (*56*). Following model estimation, the resulting beta images were normalized to MNI space using individual deformation fields obtained during preprocessing and smoothed with a 6-mm full-width at half-maximum (FWHM) Gaussian kernel.

#### Second-level analyses

The normalized and smoothed beta images generated at the first level were used for group-level random effects analysis. To this end, second level factorial GLMs were constructed using beta images corresponding to all events modeled at the first level. For significance-testing, we employed a region-of-interest (ROI) approach, based on preregistered anatomical targets: the rostral anterior cingulate cortex (rACC) and the periaqueductal gray (PAG). We derived binary masks for the right and left ACC from the Neuromorphometrics atlas (labels: ‘Left ACgG anterior cingulate gyrus’, ‘Right ACgG anterior cingulate gyrus’; http://www.neuromorphometrics.com). The PAG mask was obtained from the brainstem navigator atlas (*75*). We generated a combined mask image of these ROIs which we used for small-volume-corrected (SVC) hypothesis-testing to control for type-I error inflation. For exploration of signal outside of our preregistered regions of interest, we applied whole-brain FWE-correction. Results were considered statistically significant at *p* < .05 after correction for multiple comparisons.

To examine functional connectivity between the rACC and the PAG during pain modulation, we performed generalized psychophysiological interaction analyses (gPPI) (*76*, *77*). Seeds were defined as the subject-specific peak activation within a 6-mm-radius sphere around the group-level peak activations observed in rACC with the contrast Accepted > Declined in the SAL condition. PPI regressors were computed as the element-wise product of the extracted time series and z-scored psychological regressors forming the contrast. These, along with the psychological regressors, seed time series, and nuisance regressors, were included in new subject-level GLMs. After model estimation, the resulting beta images were taken to the second level, where contrasts of interest were computed between the PPI regressors. Correction for multiple comparisons in this connectivity analysis was restricted to the PAG ROI, the sole preregistered target for this analysis. This type of gPPI analysis was also used in exploratory connectivity analyses.

## Acknowledgments

We thank Helene Rittmeister for naloxone preparation during data collection; Darius Zokai, Christian Sprenger, Lara Hille, and Hanna Braaß for medical supervision of the pharmacological intervention. We are grateful to Janne Nold for help with planning the naloxone intervention, and to Björn Horing, Lieven Schenk, and Alexandra Tinnermann for helpful comments during study conceptualization and analysis. We further thank Katrin Bergholz, Waldemar Schwarz, and Kathrin Wendt for radiographic assistance during MR data collection, and Jürgen Finsterbusch for providing the MR sequence. The authors used a large language model (GPT-5, OpenAI) to assist with code formatting and English language editing during manuscript preparation.

## Funding

Deutsche Forschungsgemeinschaft (DFG, German Research Foundation) – Project number 449640848 (LA, CB)

European Research Council (ERC) – ERC-AdG-883892-PainPersist (LK, CB)

German Federal Ministry of Education and Research (BMBF) (OG)

Deutsche Forschungsgemeinschaft (DFG, German Research Foundation), Project number 422744262–TRR 289 (JJ, CB)

## Author contributions

Conceptualization: CB, LA

Methodology: LA, CB, OG, LK

Formal analysis: LA, CB, OG, LK

Investigation: LA, JJ

Visualization: LA, LK

Funding acquisition: CB

Project administration: CB, LA

Supervision: CB

Writing – original draft: LA

Writing – review & editing: CB, LA, OG, LK, JJ

## Competing interests

The authors declare that they have no competing interests.

## Data and code availability

All data required to reproduce the analyses and figures are publicly available at https://gin.g-node.org/asankleo/avapp_data. All code used for data processing, statistical analyses, and figure generation is available publicly at https://gin.g-node.org/asankleo/avapp_code.

## Notes

https://gin.g-node.org/asankleo/avapp_data

https://gin.g-node.org/asankleo/avapp_code

### Competing Interest Statement

The authors have declared no competing interest.

