## Supplementary Information for "The pursuit of motivational goals reduces pain through an opioidergic mechanism"

### Supplementary Text 1: Control analyses

*Control analysis I: influence of prior pain intensity on pain*

Because the painful stimuli following accepted and declined offers differed in the intensity of the preceding stimulus (80 VAS for declined, 50 VAS for accepted offers), we sought to confirm that the observed difference in Pain B perception reflected decision-related processes rather than physical carry-over effects from the preceding stimulus. To this end, we conducted a control analysis using Pain A ratings as the outcome variable. Specifically, we focused on trials with a 50-VAS Pain A stimulus that had been preceded by either an 80- or 50-VAS Pain B stimulus in the previous trial. The experiment was designed such that the inter-stimulus intervals between pain events were, on average, identical both within trials (Pain A → Pain B) and across trials (Pain $B_{t-1}$ → Pain $A_t$). This ensured that the comparison was equivalent in its physical timing properties to the main decision-related contrast. Using this subset of trials, we applied the same model specification procedure as in the primary Pain A analysis (see Methods), except that the intensity factor was omitted (as only moderate-intensity trials were included) and the intensity of the previous trial was added as a predictor.

The resulting model was specified as follows:

$$rating\ A \sim intensity\ B_{t-1} + \ decision * drug + runtrial + sesstrial + session$$

$$+ offer + \ (1 + intensity\ B_{t-1} + runtrial + sesstrial \mid \ subject)$$

Where intensity $B_{t-1}$ is the intensity of the second painful stimulus of the previous trial (either 80 VAS or 50 VAS). Results are reported in the main text. To formally test for the absence of potential carry-over effects, we conducted a Bayes factor analysis. The model described above plus a reduced model excluding the *intensity* $B_{t-1}$ factor were re-estimated using Bayesian hierarchical modeling implemented in brms *(78)* with an rstan *(79)* backend. Weakly informative priors were specified for all parameters (fixed effects: Normal(0, 5); intercept: Student-t(3, 50, 20); group-level standard deviations: Exponential(0.2); residual standard deviation: Exponential(0.1)). Posterior estimation used Markov Chain Monte Carlo (MCMC) sampling, applying four Markov chains with 4000 iterations each. Both the full and the reduced model showed excellent convergence ($\hat{R} \leq 1.01$ for all parameters), with high bulk and tail effective sample sizes and no evidence of sampling pathologies, indicating robust posterior estimation. Evidence for the presence of a carry-over effect was then assessed by comparing the full model including *intensity* $B_{t-1}$ to the reduced model excluding this predictor using bridge sampling to compute the Bayes factor.

*Control analysis II: Pain B analysis restricted to non-effort trials*

Because we observed a significant effect of the effort task on Pain B ratings, and because the effort factor was fully nested within "accept" trials, we sought to make sure that the observed decisional effect was not dependent on the execution of the physical task alone. We thus repeated the analysis of Pain B ratings described in Materials and Methods after excluding all trials in which the effort task had to be performed. The resulting model consequently did not entail the effort factor and was thus specified as follows:

$$rating\ B \sim decision * drug * rating\ A_z + runtrial + sesstrial + session + offer$$
$$+\ (1 + decision + rating\ A_z + runtrial + sesstrial \mid subject)$$

After excluding all effort trials, both the main effect of decision ($F$(1, 51.2) = 11.67, $p$ = .001) and the decision-by-drug interaction ($F$(1, 2928.42) = 5.97, $p$ = .015) remained significant (fig. S3). Consistent with the primary analysis, the decisional effect was larger in the SAL condition (mean difference = 5.07 VAS, *SE* = 1.34, $t(66.6) = 3.80$, $p = 3.22 \times 10^{-4}$) than in the NLX condition (mean difference = 2.84 VAS, *SE* = 1.32, $t(62.9) = 2.153$, $p = .035$).

**Supplementary Text 2: Reward anticipation control experiment**

This experiment was designed to isolate the effects of reward anticipation on pain perception while removing self-generated motivational choice and effort-related processes that were present in the main experiment.

*Participants*

A total of 18 healthy volunteers participated in the study (mean age = 25.83 years, *SD* = 6.15). Participants were screened using the same exclusion criteria as in the main study (Supplementary Table 3), with the exception of MRI-related exclusion criteria.

*Experimental Procedure*

Before the experiment, participants received standardized task instructions and provided written informed consent. Noxious heat stimuli were then calibrated using the same procedure as in the main experiment to determine individual temperatures corresponding to subjective pain intensities of 35, 50 and 65 VAS. Noxious heat delivery and behavioral data acquisition were otherwise identical to the main experiment, except that testing was conducted in a behavioral laboratory rather than an MRI scanner and responses were recorded using a standard computer keyboard. After calibration, participants performed the experimental task, and received monetary compensation upon completion of the session.

*Task*

The experiment employed a fully crossed 3 × 3 within-subject design with the factors cue (neutral, reward, or loss) and stimulus intensity (low, 35 VAS; moderate, 50 VAS; high, 65 VAS). Each trial began with a 2-s presentation of a monetary cue indicating either a reward, a loss or a neutral outcome (€0.00). Reward cues were preceded by a plus sign (+), whereas loss cues were preceded by a minus sign (−). Neutral cues were not preceded by a sign. To ensure attention to the monetary cues, two catch trials were interspersed in each block. On these trials, the sign preceding the monetary amount (+ or −) was replaced by a cross, and participants were instructed to press the Enter key during the 2-s cue presentation. Failure to respond on more than two catch trials would have resulted in exclusion from the experiment; however, no participant met this criterion. Reward and loss amounts were randomly sampled from a normal distribution centered on €1.50 (*SD* = €0.20). Following cue presentation, participants received a 7-s noxious heat stimulus of low (35 VAS), moderate (50 VAS) or high (65 VAS) intensity. Participants then rated perceived pain intensity during a 4-s response period using a 0-100 visual analog scale, as in the main experiment. The trial concluded with outcome presentation. Importantly, outcomes were delivered with 100% certainty on all trials. Participants completed 135 trials distributed across five blocks of 27 trials each. Cue and stimulus-intensity combinations were randomized and counterbalanced within blocks. At the end of the experiment, 10 trials were randomly selected for payment. Rewards and losses from these trials were counted toward participants' reimbursement, while a minimum payout equivalent to the hourly reimbursement amount and a maximum of €10 extra was ensured. Total duration of the experiment was approximately 1.5 hours.

*Data analysis*

Data were preprocessed using the same pipeline as the main experiment. Specifically, trials with pain ratings of 0 or 100 were excluded, as were extreme rating outliers, defined as values exceeding three times the interquartile range within each participant and stimulus intensity level.

To assess the effects of reward anticipation on pain perception, we fitted linear mixed-effects models with cue value (reward, neutral, loss) and stimulus intensity (35, 50, 65 VAS) as categorical fixed effects, including their interaction. Trial number within run and trial number across the session were included as covariates. Following the approach used in the main experiment, the maximal random-effects structure that did not produce convergence or singularity issues was determined through stepwise model building using the buildmer package *(66)*. The final model was specified as:

$$rating \sim cue * intensity + runtrial + sesstrial + \ (1 + runtrial \mid subject)$$

To quantify evidence for the absence of cue-related effects, we conducted Bayes factor analyses using the same procedure and prior specifications as in Control Analysis I. Here, we compared the full model with a reduced model excluding both the main effect of cue and all cue-related interaction terms. Full and reduced models showed excellent convergence ($\hat{R} = 1.00$ for all parameters) and high bulk and tail effective sample sizes.

*Results*

This experiment was designed to examine whether reward anticipation is sufficient to modulate pain in the absence of motivational choice and action. To establish whether the sample size was adequate to detect effects of a magnitude comparable to the decision effects observed in the main experiment, we first conducted a post-hoc sensitivity analysis. We estimated statistical power using simulation-based analyses implemented in the simr *(80)* package. Based on the observed variance structure, we simulated datasets across a range of effect sizes for the effect of a reward cue relative to a neutral cue on perception of a 50 VAS stimulus (1000 simulations per effect size) and determined the corresponding power at a sample size of $n = 18$ (Supplementary Fig. S9). These analyses indicated that in this experiment, a sample size of 18 provided 90% power ($\alpha = 0.05$, two-tailed) to detect an effect size of $d = 0.29$. Thus, the design had high sensitivity to detect cue-related effects substantially smaller than the decision-related hypoalgesic effect observed under saline in the main experiment ($d = 0.49$).

We found no evidence that reward-predictive cues influenced pain perception (main effect of cue: $F(2, 2225.75) = 1.02$, $p = .362$). Pairwise comparisons revealed no differences between reward and neutral cues (mean increase from neutral to reward: 1.20 VAS, $t(2226) = 1.41$, $SE = 0.85$, $p = .338$) or between reward and loss cues (mean increase from loss to reward: 0.43 VAS, $t(2224) = 0.51$, $SE = 0.85$, $p = .867$; Supplementary Fig. S6). Furthermore, we observed no cue-by-intensity interaction ($F(4, 2227.41) = 1.09$, $p = .361$), and no cue-related comparisons reached significance within any stimulus-intensity level (Supplementary Table 4).

Bayesian model comparison provided further evidence for the absence of cue-related effects. A reduced model excluding cue and cue-related interaction terms was approximately 38 times more likely to generate the observed data than the full model ($BF_{01} = 38.36$), indicating strong evidence in favor of the null hypothesis.

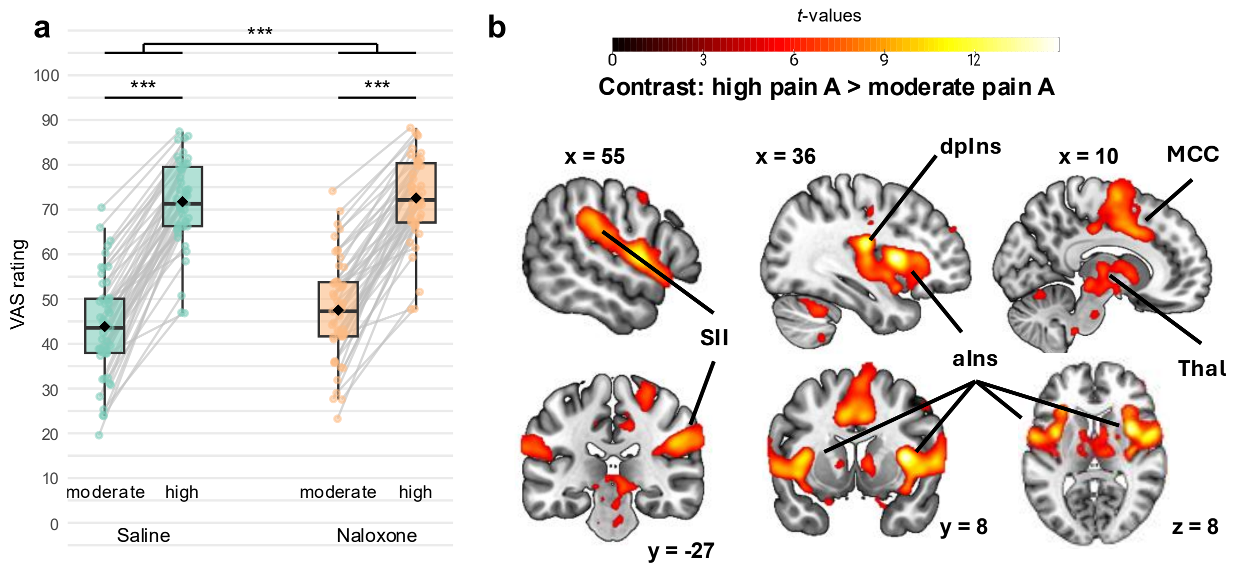


**Supplementary Fig. S1. Behavioral and neural indices of effective heat pain application. (a)** High-intensity stimuli elicited greater pain ratings than moderate-intensity stimuli for Pain A (before the decision). Naloxone (NLX) enhanced pain, with a larger effect at moderate intensity. Diamonds indicate marginal means; horizontal lines, medians; points, participant-level estimated marginal means (including random effects). $N = 49$. VAS, visual analog scale. ***$p < .001$. **(b)** BOLD activation for the contrast high Pain A > moderate Pain A was observed in typical pain-responsive regions, including secondary somatosensory cortex (SII), dorsal posterior insula (dpIns), anterior insula (aIns), mid-cingulate cortex (MCC), and thalamus (Thal). Activations are displayed at a threshold of $p < .05$ whole-brain FWE-corrected. $N = 45$.

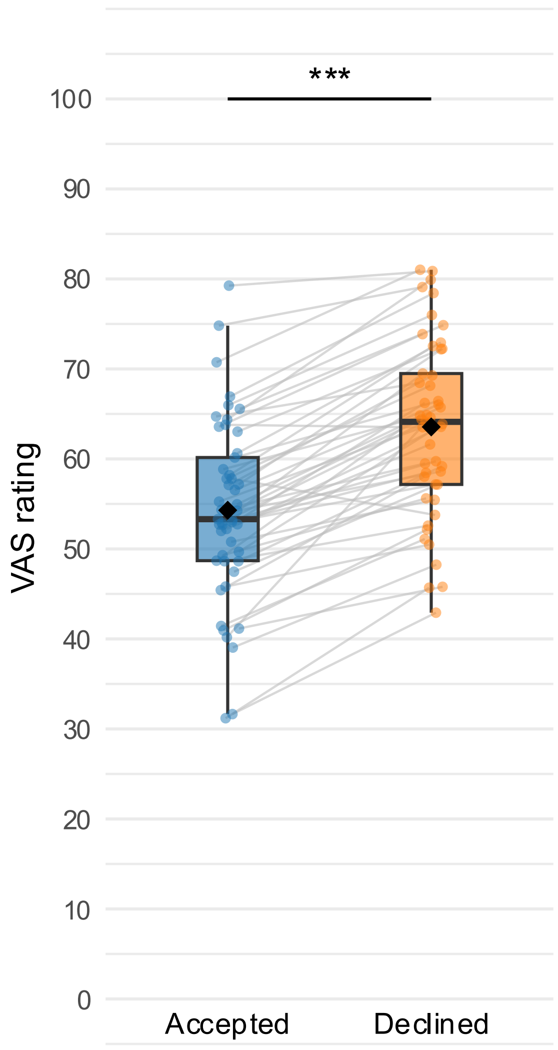


**Supplementary Fig. S2. Estimated marginal means for Pain A ratings, by subsequent decision.** Stimuli rated as more painful were more likely to be followed by declining as compared to accepting an offer, across intensities and drug conditions. Diamonds represent estimated marginal means, lines denote medians, and points represent participant-level estimated marginal means (including random effects). $N = 49$. VAS, visual analog scale. $^{***}p < .001$.

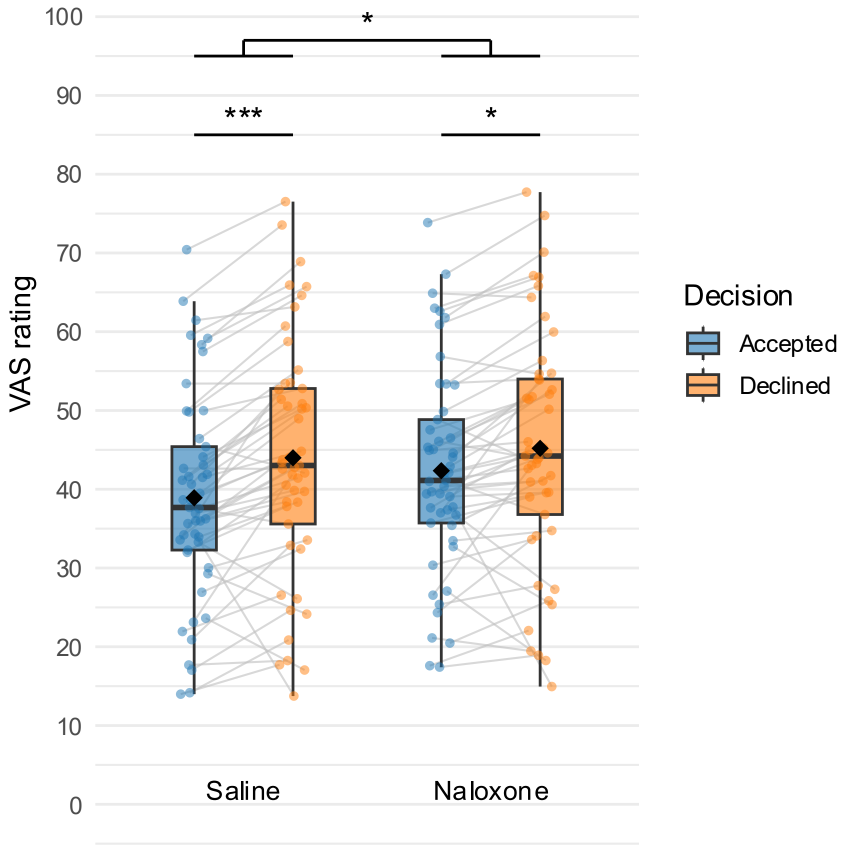


**Supplementary Fig. S3. Drug-by-decision interaction on Pain B ratings in trials without effort.** Restricting the analysis to trials in which no effort task was performed yielded the same interaction pattern observed in the full dataset. Pain B ratings were reduced following accept relative to decline decisions, and this decision-related modulation was reduced under naloxone compared with saline. This result for effort-free trials indicates that the drug-by-decision interaction is not driven by trials involving physical effort. Diamonds indicate estimated marginal means, lines denote medians, individual data points denote participant-level estimated marginal means (including random effects). $N = 49$. VAS, visual analog scale. *$p < .05$, ***$p < .001$.

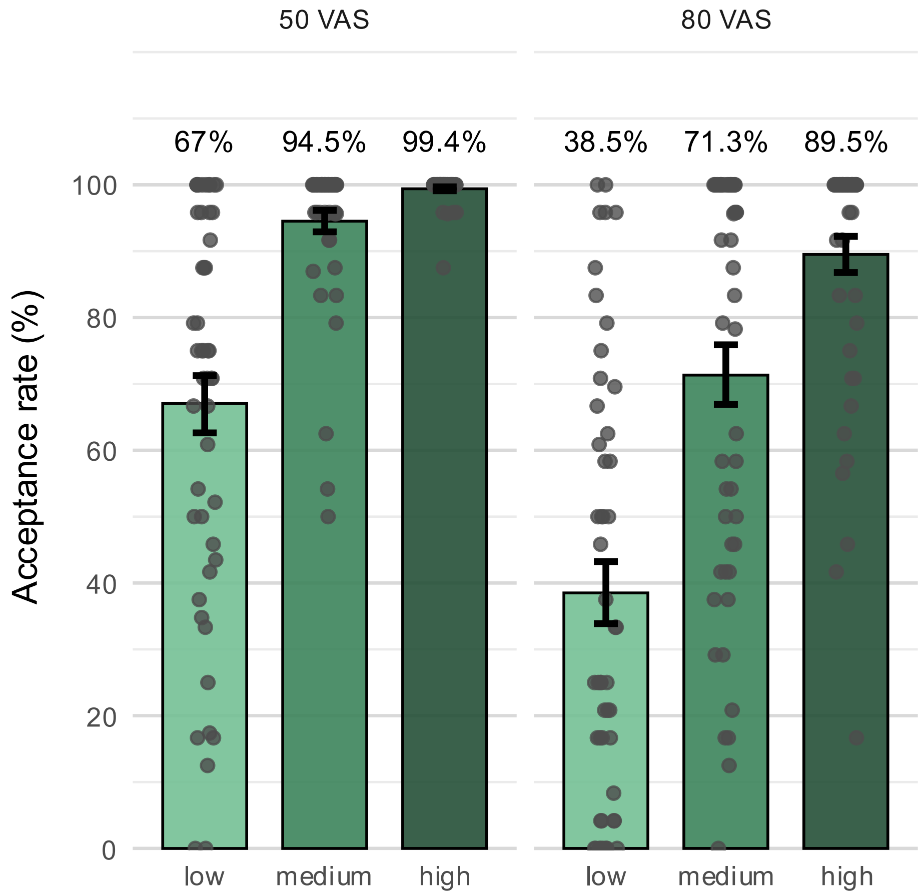


**Supplementary Fig. S4. Acceptance rates in the motivation-decision task, split by offer and Pain A intensity.** Bars represent sample means. Gray points indicate individual acceptance rates. Error bars represent standard errors of the mean. $N = 49$. VAS, visual analog scale.

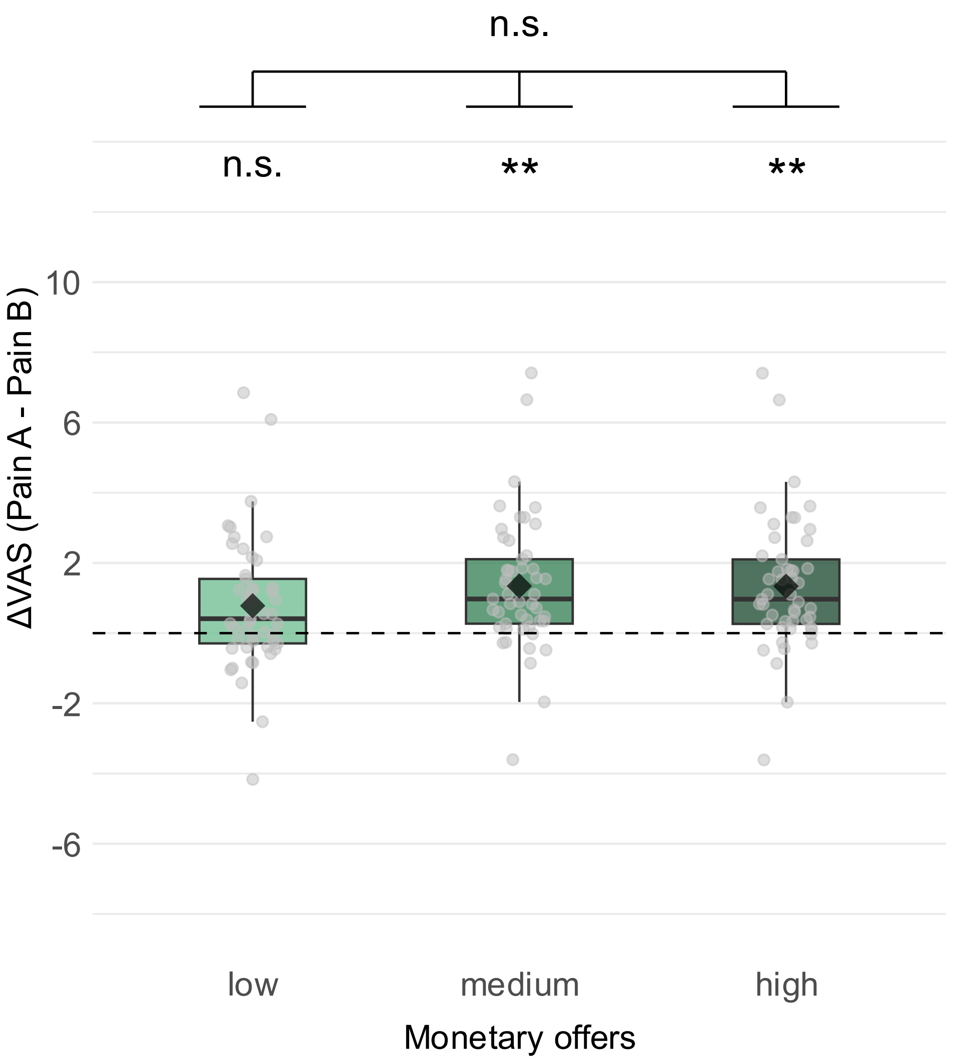


**Supplementary Fig. S5. Pain Reduction within trials, by offer.** Pain reduction within a trial was quantified as the difference between Pain A and Pain B after accepting an offer. High values indicate increased pain reduction. Sidak-corrected one-sample t-tests showed reductions significantly different from zero across drug conditions for the medium and high, but not the low offers. However, there was no global main effect of offer, indicating that pain reduction did not significantly scale with the offer amount. Diamonds indicate estimated marginal means, lines denote medians, individual data points denote participant-level estimated marginal means (including random effects). $N = 49$. VAS, visual analog scale. *$p < .05$, **$p < .01$, n.s., not significant.

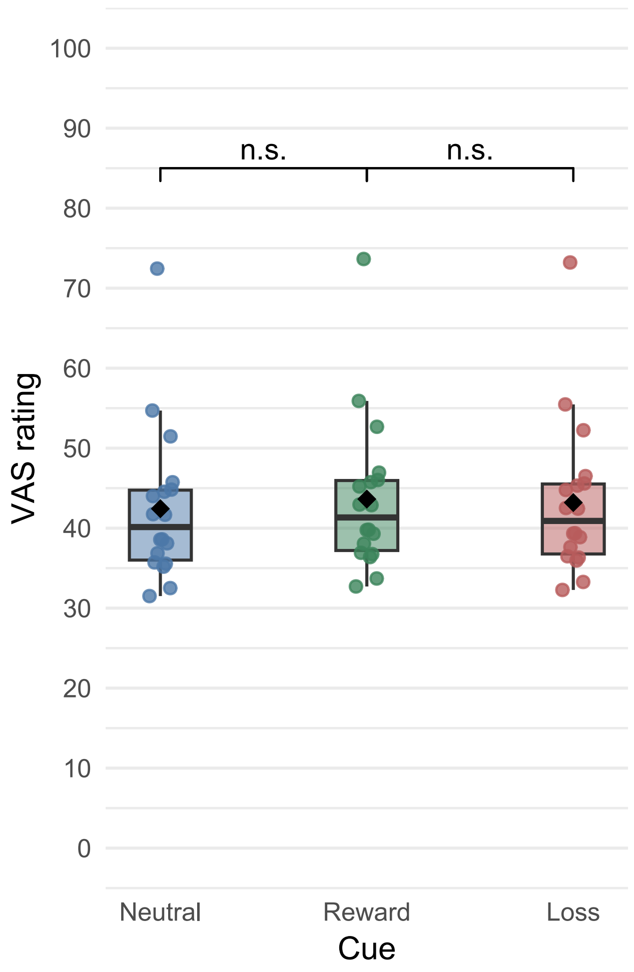


**Supplementary Fig. S6. Cue effects of the control experiment.** Estimated marginal means for ratings across pain intensities, displayed by offer cue. At 100% outcome probability and without a decisional component or an effort task, no differences were observed between the neutral outcome cue and the reward or loss cue, indicating that reward anticipation alone is unlikely to account for pain modulation. Diamonds indicate estimated marginal means; lines denote medians; points show participant-level estimated marginal means (including random effects). *N* = 18. VAS, visual analog scale. n.s., non-significant.

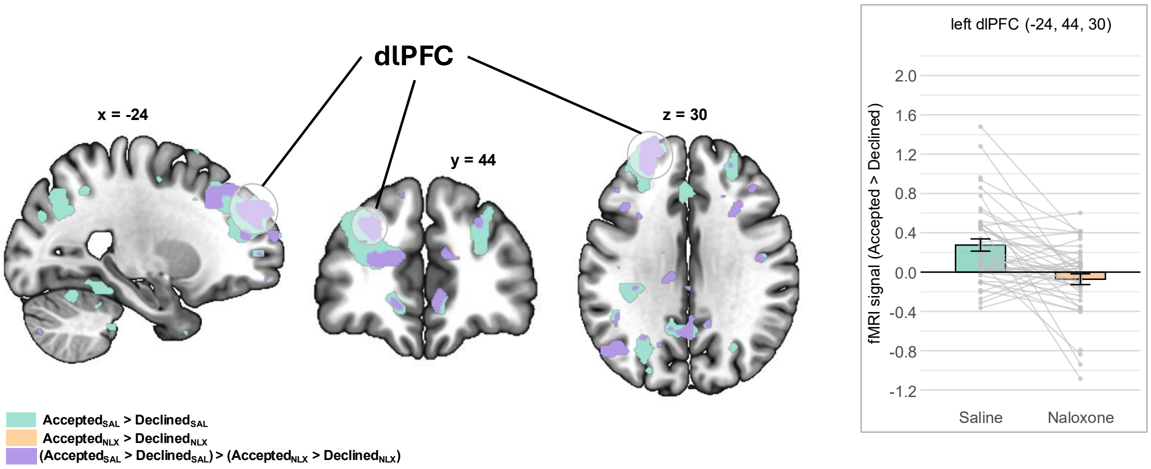


**Supplementary Fig. S7. BOLD responses during motivational pain modulation in dlPFC.** The left dlPFC showed increased signal during pain after accepting relative to declining an offer under saline, and indications of response attenuation under naloxone. Points and lines represent individual subject values. Error bars represent standard errors of the mean. Statistical maps are displayed at $p < .001$ uncorrected (for visualization only). $N = 45$. dlPFC, dorsolateral prefrontal cortex; SAL, saline; NLX, naloxone.

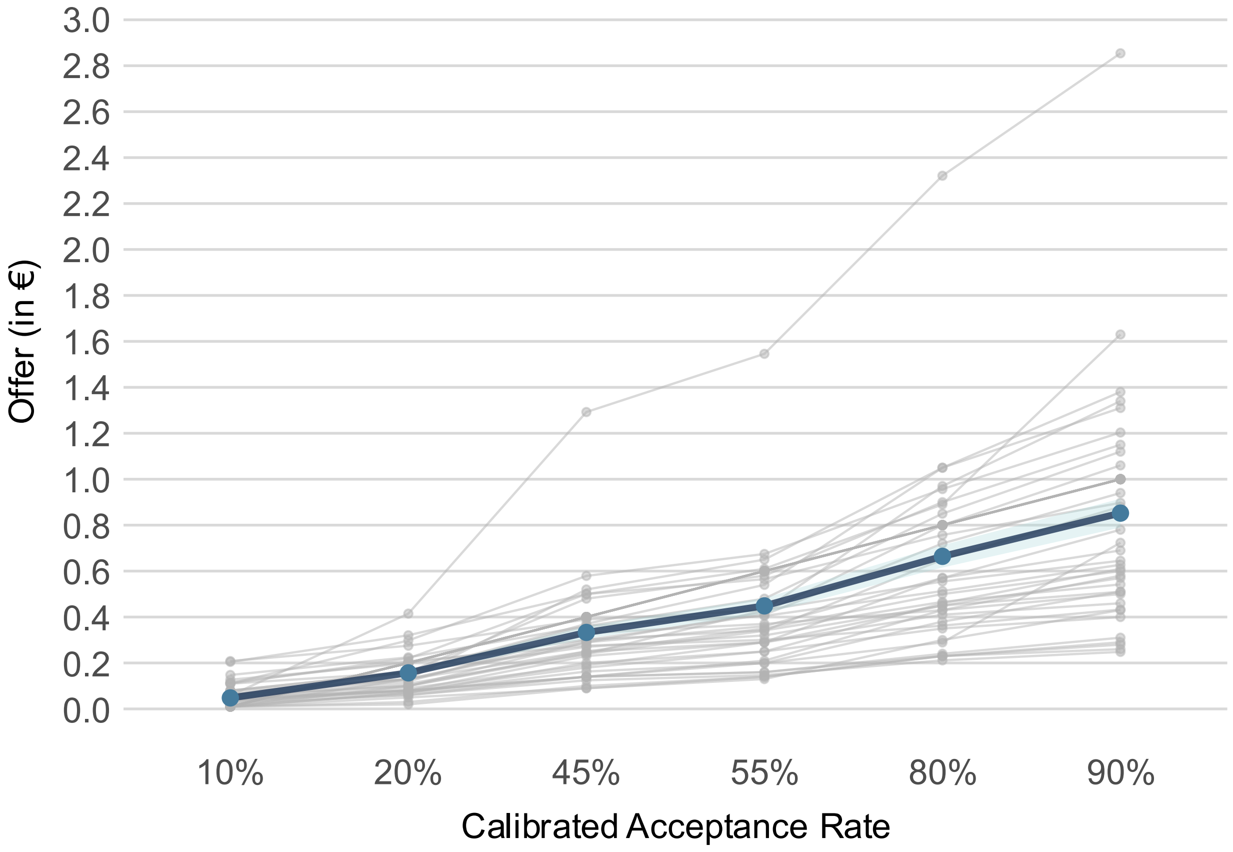


**Supplementary Fig. S8. Average and individual calibrated offer levels.** In the pre-experimental calibration session, offer levels corresponding to low (10-20%), medium (45-55%) and high (80-90%) acceptance rates were determined via logistic regression using data from the monetary calibration task. Blue line indicates sample average. Gray lines indicate individual calibration results. Blue shaded area represents standard errors of the mean. $N = 49$.

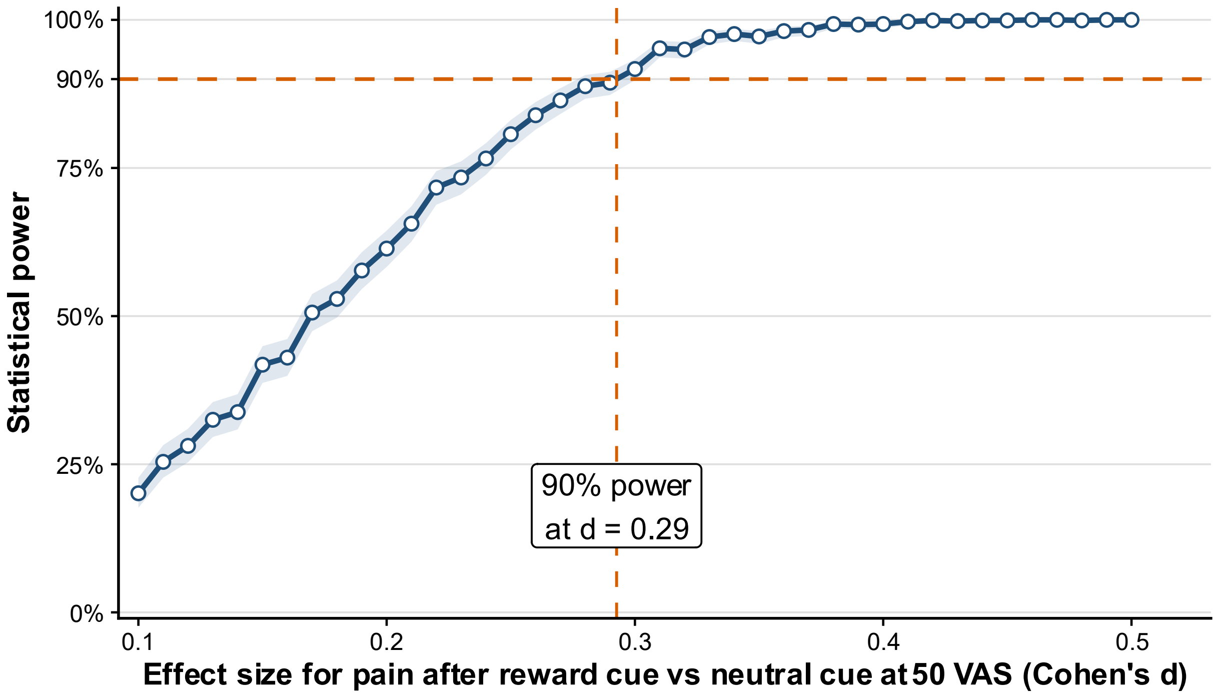


**Supplementary Fig. S9. Sensitivity power analysis for the reward anticipation control experiment.** Statistical power to detect an effect of reward anticipation on pain ratings at n = 18 as a function of standardized effect size (Cohen's d) for the contrast between pain after reward vs. neutral cues during 50 VAS pain stimulation. Power estimates were obtained using simulation-based analyses based on the fitted mixed-effects model and observed variance structure of the control experiment (n = 18; 1000 simulations per effect size). The shaded region indicates 95% confidence intervals around the estimated power. Dashed lines indicate the 90% power criterion and the corresponding detectable effect size (d = 0.29).

**Supplementary Table 1.**

Brain regions showing a significant main effect of naloxone during Pain A.

| Region | MNI (x, y, z) | *Z* | *p* |
|---|---|---|---|
| **Cerebellum** | | | |
| Lobule VI right | 30, -60, -21 | 4.90 | .041 |
| Lobule VI left | -26, -60, -20 | 5.57 | .002 |
| Crus I right | 44, -50, -28 | 5.60 | .001 |
| Crus II left | -22, -90, -33 | 5.33 | .005 |
| **Thalamus** | | | |
| left anterior | -2, -4, 6 | 5.32 | .006 |
| right ventral anterior | 8, -15, -12 | 4.87 | .046 |

**Note.** Peak coordinates are reported in MNI space. Listed regions correspond to local maxima within clusters showing a significant main effect of naloxone during Pain A for the contrast $[(\text{Accepted}_{NLX} + \text{Declined}_{NLX}) > (\text{Accepted}_{SAL} + \text{Declined}_{SAL})]$ at $p < .05$ whole-brain FWE-corrected. *Z*-values indicate peak voxel statistics.

**Supplementary Table 2.**
Individual reasons for study dropouts.

| ID | Reason for exclusion |
|---|---|
| 3 | Unpleasant side effects |
| 8 | Potential ECG abnormalities |
| 11 | Technical difficulties |
| 13 | Pain became intolerable |
| 19 | Claustrophobia during scanning |
| 20 | IV access issues |
| 28 | IV access issues |
| 29 | Withdrew before second session due to medical concerns |
| 37 | Withdrew spontaneously |
| 41 | Withdrew before second session due to time constraints |
| 46 | IV access issues |
| 50 | Failed MRI safety screening |
| 59 | Participated in another medical study in the preceding week |
| 67 | Took codeine before experimental session |

**Supplementary Table 3.**

Preregistered inclusion and exclusion criteria.

| Inclusion and exclusion criteria | |
|---|---|
| **Inclusion criteria** | Healthy, adult participants |
| | Native-level proficiency in German |
| | Age limit: 18–45 years |
| | Right-handedness |
| **Exclusion criteria** | Acute or chronic pain (including soreness) |
| | Presence of acute or chronic somatic or psychiatric illness (based on self-report) |
| | MRI-specific exclusion criteria (e.g., claustrophobia, pacemaker, non-MR-compatible metallic implants) |
| | Participation in studies involving medication, or regular use of medication (except thyroid, allergy medication, occasional use of painkillers, or contraceptives) within the past 2 months |
| | Use of painkillers within 24 hours prior to the examination |
| | Fear of needles or blood sampling |
| | Pregnancy or breastfeeding |
| | Acute skin disease or injury in the stimulation area |
| | Acute illness or symptoms of the respiratory tract (e.g., exercise-induced asthma) |
| | Past or current physical opioid dependence |
| | Cardiovascular disease (e.g., angina pectoris) |
| | Neurological disease |
| | Immune system disorder |
| | Gastrointestinal disease |
| | Use of cardiotoxic substances (e.g., cocaine, methamphetamine, cyclic antidepressants, calcium antagonists, beta blockers, digoxin) |
| | Metallic implants in the body (e.g., intrauterine devices, stents, bone screws) |
| | Electromagnetic or electric implants (e.g., pacemakers) |

**Supplementary Table 4. Pairwise comparisons for cue effects within the three intensity levels.**

| Comparison | Estimate | *SE* | df | *t* | *p* |
|---|---|---|---|---|---|
| **35 VAS** | | | | | |
| Reward vs. neutral | 0.93 | 1.53 | 2225 | 0.61 | .814 |
| Reward vs. loss | 1.02 | 1.53 | 2228 | 0.66 | .785 |
| Loss vs. neutral | -0.08 | 1.51 | 2226 | -0.06 | .998 |
| **50 VAS** | | | | | |
| Reward vs. neutral | 0.44 | 1.45 | 2226 | 0.30 | .952 |
| Reward vs. loss | 1.26 | 1.44 | 2226 | 0.87 | .659 |
| Loss vs. neutral | -0.82 | 1.45 | 2226 | -0.57 | .838 |
| **65 VAS** | | | | | |
| Reward vs. neutral | 2.23 | 1.45 | 2225 | 1.54 | .274 |
| Reward vs. loss | -0.97 | 1.46 | 2229 | -0.67 | .784 |
| Loss vs. neutral | 3.21 | 1.46 | 2230 | 2.20 | .072 |

**Note.** Pairwise comparisons were computed from estimated marginal means obtained from the linear mixed-effects model described in Supplementary Text 2. Positive estimates indicate higher pain ratings for the first cue condition relative to the second. *P* values were Tukey-corrected within each stimulus intensity level. VAS, visual analog scale. $N = 18$.
